# Correlating High-dimensional longitudinal microbial features with time-varying outcomes with FLORAL

**DOI:** 10.1101/2025.02.17.638558

**Authors:** Teng Fei, Victoria Donovan, Tyler Funnell, Mirae Baichoo, Nicholas R. Waters, Jenny Paredes, Anqi Dai, Francesca Castro, Jennifer Haber, Ana Gradissimo, Sandeep S. Raj, Alexander M. Lesokhin, Urvi A. Shah, Marcel R. M. van den Brink, Jonathan U. Peled

## Abstract

Correlating time-dependent patient characteristics and matched microbiome samples can identify biomarkers in longitudinal microbiome studies. Existing approaches typically repeat a pre-specified modeling approach for all taxonomic features, followed by a multiple testing adjustment step for false discovery rate (FDR) control. In this work, we develop an alternative strategy of using log-ratio penalized generalized estimating equations, which directly models the longitudinal patient characteristic of interest as the outcome variable and treats microbial features as high-dimensional compositional covariates. A cross validation procedure is developed for variable selection and model selection among different working correlation structures. In extensive simulations, the proposed method achieved superior sensitivity over the state-of-the-art methods with robustly controlled FDR. In the analyses of correlating longitudinal dietary intake, bloodstream infection status, and microbial features from matched samples of cancer patients, the proposed method effectively identified gut health indicators and clinically relevant microbial markers, showing robust utilities in real-world applications. The method is implemented under the open-source R package `FLORAL`, which is available at (https://vdblab.github.io/FLORAL/).

## 1 Introduction

Longitudinal patient data collection has become increasingly more prevalent in microbiome studies, where microbial samples are paired with longitudinal patient data, such as bloodstream infection (BSI) [1, 2], dietary intake [3–5], body mass index (BMI) [6], blood counts [7], immune cell measurements [8], and metabolite abundance [9]. Rich longitudinal clinical data offer valuable opportunities to explore microbial associations with various temporal variables from the clinical side, which further assists hypothesis generation in basic biological research which can be facilitated by mouse models derived based on the clinical observations. For example, it is of interest to investigate the associations between microbial taxa and the amount of fiber intake for patients undergoing allogeneic hematopoietic cell transplantation (allo-HCT), which will contribute to identifying strategies for dietary interventions.

Despite the availability of longitudinal microbiome data, the relevant literature in statistical and computational methods is limited. Feature selection methods have been proposed to correlate longitudinal microbial features with a continuous, binary, or time-to-event disease outcome observed after the last time point of sample collection [10, 11], which is not generalizable to the paired longitudinal microbiome and longitudinal patient characteristic data scenario. A versatile dimension reduction method was developed to perform temporal tensor decomposition for the taxa trajectories with respect to taxa, individual, and temporal patterns [12], which demonstrated utility in correlating pre-specified patient groups and longitudinal microbial patterns as temporal loadings. Nevertheless, the method did not provide an explicit feature selection approach which incorporates dynamically changing patient characteristics. In contrast to the above methods, the mixed-effect model is a more widely applied class of methods that effectively incorporates longitudinal patient characteristics into the models of individual taxon trajectories [13–15]. Typically, the same mixed-effect model structure is repeatedly applied to model all taxonomic features, followed by a false discovery rate (FDR) control procedure across all models to identify significant associations between longitudinal patient characteristics and taxa. In practice, however, different taxa may have largely varying stability across repeated observations and highly variable patient-specific distributions, making it challenging to apply a single model configuration (for example, linear time effect and random intercept) for hundreds of taxa. Additionally, the sparse and compositional nature of microbiome data brings additional challenges in modeling the abundance trajectories of taxa, which motivated the use of complex modeling strategies to account for zero-inflation, over-dispersion and potentially non-linear associations [15]. As a common alternative to the mixed-effect model, generalized estimating equations (GEE) have also been applied to study microbial associations with longitudinal characteristics [16], yet the method focused on inferring on global microbial associations instead of taxon-level associations.

In this work, we propose a penalized log-ratio GEE model to select longitudinal microbial features associated with a longitudinal patient outcome. Here, the *outcome* variable refers to any patient characteristics collected at roughly the same time as each microbiome sample, such as dietary intake, BMI, CD4 T-cell count from flow cytometry, or the concentration of a certain short-chain fatty acid from a metabolic assay (**Fig.1A**). Unlike the widely applied mixed-effect models, we treat the longitudinal patient characteristic as the outcome variable and the microbial taxa as covariates in a multivariable regression framework. As shown in **Fig.1B**, the proposed GEE model accounts for the within-subject dependency of the outcome variable with common working correlation structures such as independence, compound symmetry, and autoregressive (AR)-1 structures. Similar to the “fitting a log-ratio lasso” (`FLORAL`) regression framework we previously developed [11], we assume a sparse set of taxa are associated with the longitudinal outcome, where the penalized estimating procedure is extended from the standard approaches [17, 18] by adding the zero-sum constraint to account for the compositional nature of the covariates. We develop a model and variable selection procedure based on the cross-validated deviance residual [19] with two-step feature filtering to further control the false discovery rate (FDR) [11, 20]. The method is publicly available as a new module within the R package `FLORAL`.

**Figure 1:**
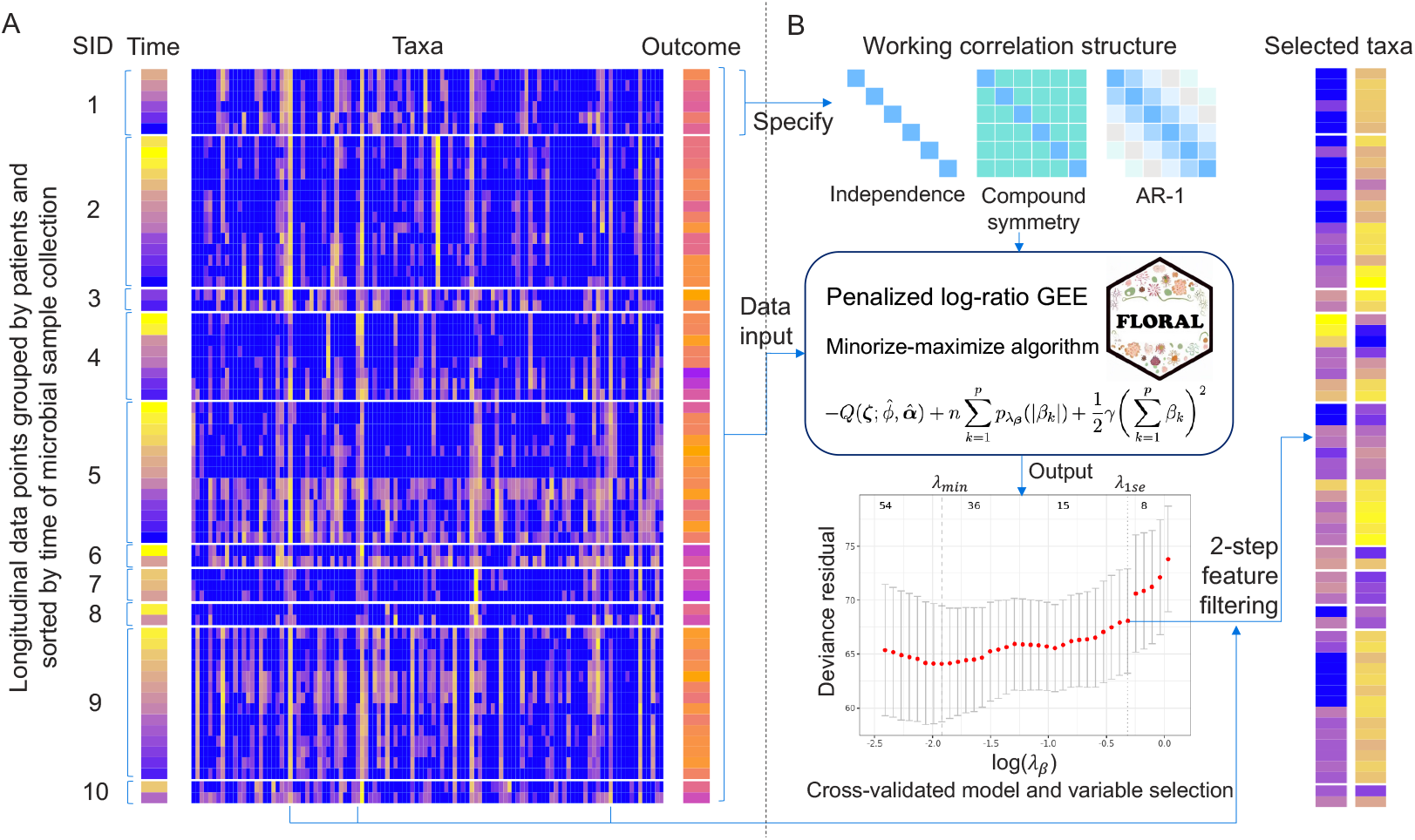
Flow chart of using the proposed penalized log-ratio generalized estimating equation (GEE) approach. **A.** Heatmaps of the input data based on 10 randomly chosen patients from the MSK allo-HCT cohort, including longitudinal taxa features, the corresponding longitudinal outcome variable (fiber intake), and time of sample collection, where brighter colors represent larger numerical values. Data points were grouped by artificially assigned subject IDs (SID) and sorted by time of sample collection. **B**. The pipeline of the proposed method. First, a user-specified working correlation structure is required for the GEE model. Then the penalized log-ratio GEEs are solved by a minorize-maximize algorithm with a zero-sum constraint. Finally, cross validations are used to determine penalty parameters (*λ*_min_, *λ*_1se_) based on deviance residual, where the selected features will be further screened by an additional ratio-based procedure (2-step filtering). The heatmaps of two selected taxa were shown on the right.

Compared to the mixed-effect models, the proposed penalized log-ratio GEE model addresses several challenges of modeling individual taxon trajectories by flipping the roles of longitudinal taxa and longitudinal patient characteristics in the regression framework. Instead of modeling the highly sparse, volatile, and compositional taxa features, we focus on modeling the more tractable and stable patient characteristics which can be conveniently depicted by Gaussian or binomial link functions. In addition, the log-ratio covariate space effectively transforms zero-inflated quantities into a continuous variable space, which mitigates the computational burdens caused by complex models. Moreover, the proposed approach focuses on modeling the marginal expectation of one fixed outcome variable, where the non-linear associations between the outcome variable and time can be easily captured by using splines [21] without specifying the forms of subject-specific random effects (e.g. random intercepts and random slopes). Finally, the cross-validated variable selection process of the proposed method is more data-driven than the existing methods which are based on a pre-specified threshold of significance. We demonstrate by extensive simulations that the proposed method requires smaller number of patients or samples to achieve similar variable selection performances as the mixed-effect models while controlling for FDR. In real-data analyses, the proposed method identifies meaningful associations between fiber intake or BSI and taxa abundance from three studies conducted at Memorial Sloan Kettering Cancer Center (MSK) with different patient populations [3, 4, 22], showing strong practical utilities in detecting clinically relevant microbial markers.

## 2 Methods

### 2.1 Notations and model formulations

Let *Y*_*ij*_ denote the *j*th observation of the longitudinal outcome for the *i*th subject, *i* = 1, …, *n, j* = 1, …, *m*_*i*_. Let ***X***_*ij*_ denote the associated *p ×* 1 microbial count vector and ***W*** _*ij*_ denote the *L ×* 1 confounder feature vector.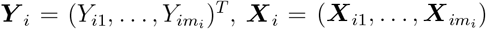, and 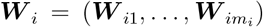. While the number of observations *m*_*i*_ varies across different subjects in practice, we assume *m*_*i*_ = *m <* ∞ without loss of generality in the following description of the proposed model.

In the proposed generalized estimating equation (GEE) model for compositional covariates, we adapt the framework of log-contrast framework [11, 23] to model the mean and the variance of the longitudinal outcome *Y*_*ij*_:

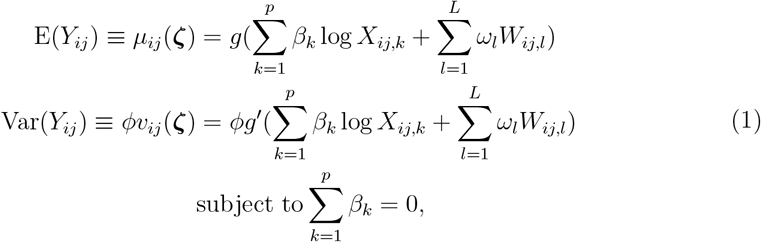

where **ζ** = (***β***^*T*^, ***ω***^*T*^)^*T*^ is the vector of unknown regression coefficients, including ***β*** = (*β*_1_, …, *β*_*p*_)^*T*^ for log-transformed compositional features and ***ω*** = (*ω*_1_, …, *ω*_*L*_)^*T*^ for non-compositional features. In addition, *g*(·) is a differentiable link function, and *ϕ* is a scaling factor which can be assumed as fixed or to be estimated. We impose a zero-sum constraint 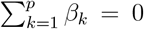 for the unknown coefficients ***β*** associated with the log-count features log(***X***_*ij*_), which makes the model equivalent to an unconstrained linear model of all possible log-ratios of the compositional features [11, 23], thus accounting for the compositional nature of the features. Similar to the GEE model [24], we also consider a working correlation structure to depict the correlations within the repeated measurements ***Y*** _*i*_ from the same individual or cluster, where the working correlation matrix of ***Y*** _*i*_ is denoted by ***R***(***α***). Popular choices of ***R***(***α***) include independence, compound symmetry (or exchangeable), or autocorrelation (AR)-1, where ***α*** follows different configurations. It then follows that 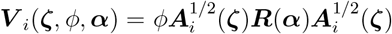 is the variance-covariance matrix of ***Y*** _*i*_, where ***A***_*i*_(**ζ**) = diag{*v*_*i*1_(**ζ**), …, *v*_*im*_(**ζ**)}. To investigate the association between longitudinal compositional features and the corresponding outcomes, the main parameter of interest is the effect size vector ***β***, while *ϕ* and ***α*** are treated as nuisance parameters in the estimation procedure. In practice, the number of compositional features *p* can be larger than the number of subjects *n* and the number of samples *n × m*. We assume that only a sparse set of the features are associated with the outcome, which means the majority of the elements in ***β*** are zeros.

### 2.2 Estimation procedure

Given the above formulation of the first two moments of ***Y***, we obtain an unbiased constrained estimating function for **ζ** given *ϕ* and ***α***

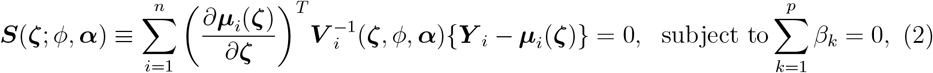

which follows the same form as the classic GEE [24] with an additional zero-sum constraint. To impose sparsity of ***β***, one natural approach is to consider a regularized regression framework such as lasso [25]. However, unlike the standard lasso regression which minimizes a penalized negative log-likelihood function, our model assumption (1) does not have an explicit likelihood function to construct an optimization problem. Instead, the proposed estimating equation (2) is a constrained zero point finding problem. Therefore, we adapt alternative strategies for penalized estimating equations [17] to fulfill the regularization of the coefficients. Specifically, we extended the penalized generalized estimating equations (PGEE) [18] framework, where we incorporated the zero-sum constraint into the PGEE, established a more systematic model and feature selection mechanism via cross-validation, and developed easily accessible software for wide applications.

The PGEE function with zero-sum constraint is formulated as

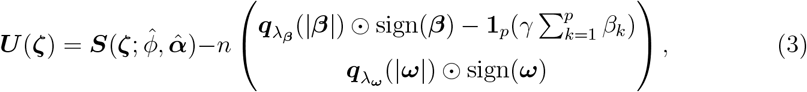

where 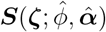 is the estimating function as defined in (2), where 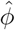 and 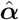 are estimates of *ϕ* and ***α*** to be updated at each iteration of the algorithm. Here, 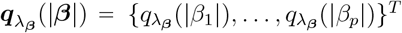 and 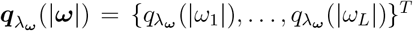 are penalty functions corresponding to each element of **ζ**, where the non-negative penalty parameters *λ*_***β***_ and *λ*_***ω***_ are separately specified for ***β*** and ***ω*** to flexibly impose penalties to compositional features while adjusted for covariates. In addition, ⊙ denotes the element-wise multiplication operator, sign(***β***) = {sign(*β*_1_), …, sign(*β*_*p*_)}^*T*^ and sign(***ω***) = {sign(*ω*_1_), …, sign(*ω*_*L*_)}^*T*^, where sign(*x*) = *I*(*x >* 0) - *I*(*x <* 0). The term 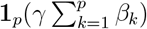 corresponds to the zero-sum constraint, where **1**_*p*_ is a *p*-vector with all elements equal to one and *γ* is the penalty parameter which governs the strength of the constraint 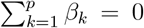. The proposed estimator 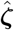 satisfies 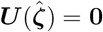.

The formulation of (3) can be derived from the following target function of an optimization problem

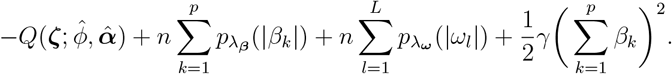

Here *Q*(**ζ**; *ϕ*, ***α***) is the quasi log-likelihood corresponding to ***S***(**ζ**; *ϕ*, ***α***), which satisfies 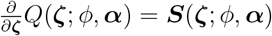. Functions 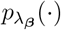 and 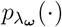 determine the form of the penalties, such as *L*_1_ or *L*_2_ penalty terms, which satisfy 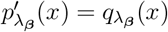 and 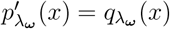. For example, *p*_*λ*_(|*x*|) = *λ*|*x*| and *q*_*λ*_(|*x*|) = *λ* for lasso penalty [25], while 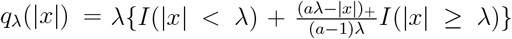 for SCAD penalty [26] with *a >* 2. The term 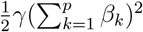 utilizes the penalty method to penalize the quantity of ∑ *β*_*k*_ with the penalty parameter *γ* [27]. It can be shown that (3) approximates the negative differentiation of the above target function with respect to **ζ**, such that we transform the optimization problem to the estimating equation solving problem of ***U*** (**ζ**) = **0**. In practice, ***β*** and ***ω*** are set to be penalized by the same type of penalty functions. In addition, *λ*_***ω***_ is set as a fraction of *λ*_***β***_ with *λ*_***ω***_ = *rλ*_***β***_, *r* ∈ [0, 1]. The penalty parameter *γ* is set as 10^5^ to numerically enforce the zero-sum constraint.

As described in [17], a minorize-maximize (MM) algorithm with local quadratic approximations for ***q***_*λ*_(| · |)sign(·) is used to obtain the estimates 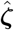. In the following, we use subscript {*i*} to denote the index of pathwise solutions with respect to a series of *λ*_***β***_, and use superscript (*j*) to denote the index of iterations of the MM algorithm with a fixed *λ*_***β***_. Like lasso regression, we obtain estimates 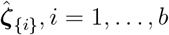 *b* along a decreasing path of *λ*_***β***{1}_, …, *λ*_***β***{*b*}_, where the largest parameter *λ*_***β***{1}_ is determined by a standard formula used for lasso regression while treating all observations independent of each other [11], while the value of the smallest parameter *λ*_***β***{*b*}_ is by default set as 0.01*λ*_***β***{1}_. The initial value 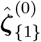 associated with *λ*_***β***{1}_ is set as zeros, while the initial value 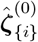 associated with *λ*_***β***{*i*}_ can be set as the “warm-start” estimates 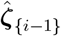 associated with *λ*_***β***{*i*−1}_. Given *λ*_***β***_ and estimate 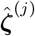 after *j*th iteration, the parameters are updated in the (*j* + 1)th iteration as following. First, let 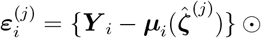 vecdiag 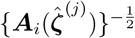 be the Pearson residual of the *i*th subject, the we update 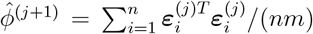 as described in Table 45.13 of [28]. Then 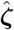 is updated as

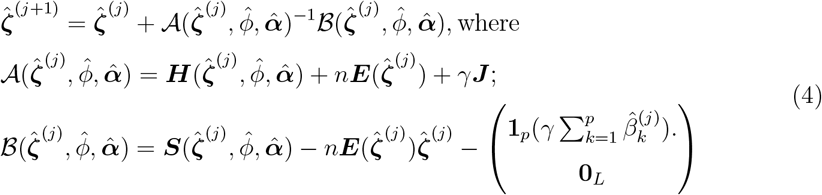

Here, ***J*** is a (*p* + *L*) *×* (*p* + *L*) matrix with the upper left *p × p* entries equal to one and zero elsewhere, **0**_*L*_ is a *L*-vector with all entries equal to zero,

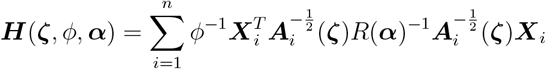

and

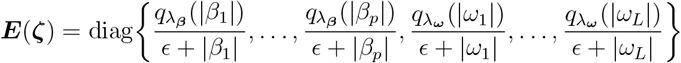

is a (*p* + *L*) *×* (*p* + *L*) diagonal matrix such that ***E***(**ζ**)**ζ** is a quadratic approximation of 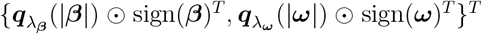 sign(***ω***)^*T*^ }^*T*^. It can be shown that that 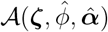 is the derivative matrix of 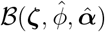 with respect to **ζ**, forming a close connection to the Newton-Raphson algorithm. In our implementation, we set *ϵ* = 10^−6^ and set the convergence criterion as ∥**ζ**^(*j*+1)^ - **ζ**^(*j*)^∥_∞_ *<* 10^−3^. After reaching convergence or exceeding 100 iterations for a given *λ*_***β***_, parameter estimates with absolute values smaller than 10^−3^ are set as zeros as suggested by [18, 29]. The algorithm described above is summarized as Algorithm 1, which is implemented in R package FLORAL with RcppArmadillo [30].

### 2.3 Variable selection via cross validation and 2-step filtering

We utilize *K*-fold cross validation to determine the values of the penalty parameters. For each choice of the penalty parameter *λ*_***β***_, we split the data into *K* folds by individual identifiers, such that all observations from a certain individual will be assigned to the same fold. Then for *k* = 1, …, *K*, the proposed model is fitted using all except the *k*th fold, then we calculate the deviance residual [19] according to the distribution family used by the GEE for all observations from the *k*th fold. The cross-validated average deviance residual is then used for model selection, where the penalty parameter achieving the smallest cross-validated deviance residual (*λ*_min_) and the largest penalty parameter with its deviance residual within one standard-error of the smallest deviance residual (*λ*_1se_) are widely used choices [11, 25]. Subsequently, we report the features with non-zero regression coefficients at *λ*_min_ and *λ*_1se_. The same cross-validation procedure is also used for model selection across different choices of working correlation structures, where the working correlation structure achieving the smallest cross-validated deviance residual is treated as the best option.

#### Algorithm 1 Iterative optimization algorithm for solving 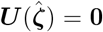 with given *λ*_***β***_ and *γ*. Note that the following algorithm assumes no intercept term. The algorithm with intercept term can be derived similarly. ⊙ denotes element-wise multiplication. Given a general square matrix **M**, the vecdiag(**M**) operator creates a vector whose elements are on the diagonal of the matrix.

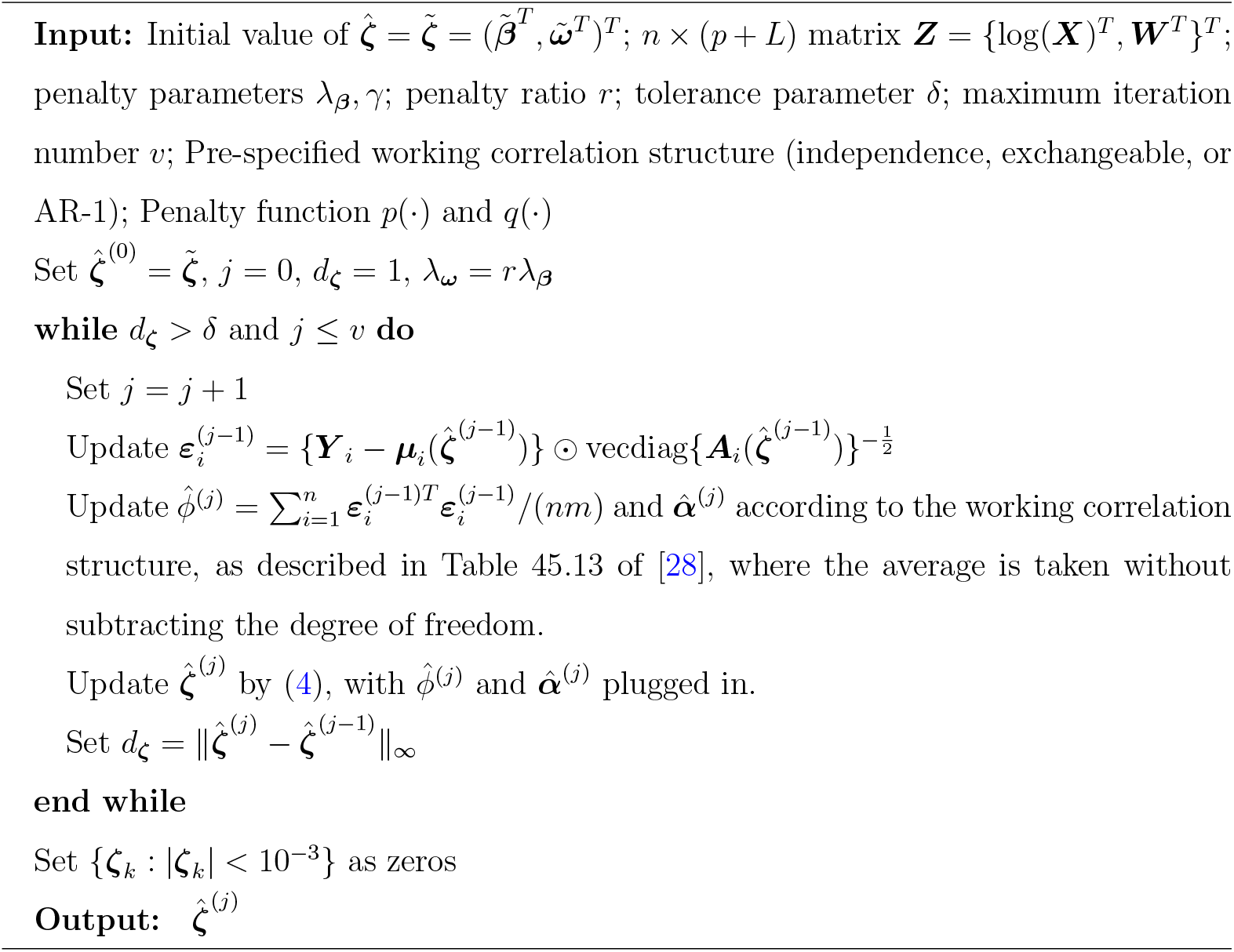

Following the cross-validated feature selection, we further implement the step-2 feature selection procedure [11, 20]. Specifically, all pairs of log-ratios based on selected features at *λ*_min_ (and *λ*_1se_) are refitted by the PGEE method without the zero-sum constraint to achieve a higher sparsity of feature selection and ratio-based model interpretation. Similar to the other regularized log-ratio regression methods implemented in the FLORAL package, the PGEE model also allows users to repeat the cross validation steps for multiple times with random fold splits, then summarize the frequency of variable selection out of all the repeats.

### 2.4 Method assessment and benchmarking

We conducted extensive simulations and real-data analysis to study and benchmark the performance of the proposed constrained PGEE method under various scenarios. Due to the scarcity of tools developed for feature selection for the longitudinal microbiome sample - longitudinal outcome data structure, we mainly focused on comparing feature selection performances within the scope of PGEE. For log-transformed data, we compared the zero-sum constrained PGEE model implemented by `FLORAL` and the standard unconstrained PGEE model. Then we also investigated the performance of PGEEs with relative abundance data and CLR-transformed data. We also applied the popular `MaAsLin2` and `MaAsLin3` packages, which use mixed-effect regression models of microbial abundance over longitudinal covariates, to better understand the pros and cons of the proposed clinical outcome-oriented modeling strategy and the popular taxa-oriented modeling strategy.

#### 2.4.1 Simulations

Let *n* be the number of individuals, *m* be the number of samples per individual, and *p* be the number of features. Longitudinal microbiome samples were simulated following a similar approach as described in [11]. We assume that the samples were observed at time *t* = 0, …, *m* - 1 for each individual. First, we simulated the true count of longitudinal microbiome data ***C*** based on a logistic-normal model [23], then the log-ratios consisting of the first ten features were utilized to generate the longitudinal outcome ***Y*** with a given correlation structure within each individual. Finally, the observable count data ***X*** was generated based on the true count data ***C*** and a randomly simulated sequencing depth for each sample. We considered scenarios with continuous outcomes and binary outcomes.

For the *i*th individual at time *t*, we first generate a vector ***x***_*i*_(*t*) = {***x***_*i*1_(*t*), …, ***x***_*ip*_(*t*)} from a *p*-variate normal distribution *N*_*p*_{***ξ***(*t*), **Σ**(*t*)} with ***ξ***(*t*) = {*ξ*_1_(*t*), …, *ξ*_*p*_(*t*)}^*T*^. We let *ξ*_*k*_(0) = log *p* for *k* = 1, 2, 3, 5, 6, 8 and otherwise *ξ*_*k*_(0) = 0, such that there are six features with higher abundance at time 0. In addition, we assume

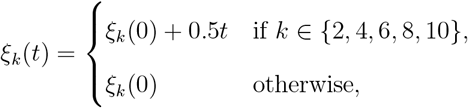

where 0.5 is the slope with respect to time for five pre-specified features. Regarding the covariance parameter **Σ**(*t*), we set the variances 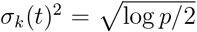 for *k* = 1, 2, 3, 5, 6, 8 and otherwise 1, such that the features with higher baseline abundances also have higher variations throughout the follow-up. We also set the covariance **Σ**_*j*,*k*_(*t*) = *ρ*^|*j*−*k*|^, *ρ* ∈ [0, 1) between features *j* and *k*. For the same individual, we also impose a sample-wise correlation of 0.4 for the first ten features. Additionally, we specify a sparsity level of 0.8 and randomly let 80% of the entries in ***x***_*i*_ to be −∞ to create zeros in compositions. After the above steps, we obtain the unobservable underlying time-dependent composition vector ***c***_*i*_(*t*) for *t* = 0, …, *m* - 1, where the *k*th entry satisfies *c*_*ik*_(*t*) = exp{*x*_*ik*_(*t*)}*/* ∑_*d*_ exp{*x*_*id*_(*t*)}. With the true composition ***c***_*i*_(*t*), we assume that the total count is 10^6^ for each sample and generate the true count vector ***C***_*i*_(*t*) from a multinomial distribution with 10^6^ counts and probability vector ***c***_*i*_(*t*).

With true count vector ***C***_*i*_(*t*), we calculate the underlying true linear predictor ***l***_*i*_ = {*l*_*i*_(0), …, *l*_*i*_(*m* - 1)}^*T*^ which consists of five log-ratios from the first ten features

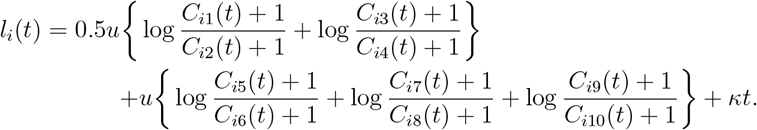

Here, a pseudo value 1 is added to all counts to make sure the log-transformations are well defined, *κ* is the constant time effect on the longitudinal outcome, and effect size *u* governs the strength between compositional features and the associated longitudinal outcome *Y*_*i*_(*t*). Given *l*_*i*_(*t*), the continuous outcome vector ***Y*** _*i*_ = {*Y*_*i*_(0), …, *Y*_*i*_(*m*−1)}^*T*^ is generated from ***Y*** _*i*_ = ***l***_*i*_ + ***ε***_*i*_, where ***ε***_*i*_ follows a zero-mean normal distribution with variance one and an exchangeable correlation 0.8 across repeated measurements. For binary outcomes, we adapt the approach from [31] to generate correlated binary outcomes from the probability vector {1 +exp{−*l*_*i*_(*t*)}}^−1^, *t* = 0, …, *m*− 1 with an exchangeable correlation 0.8 within the same individual. After generating the outcome ***Y*** _*i*_ by the underlying true count ***C***_*i*_(*t*), the observable count vector ***X***_*i*_(*t*) for each individual *i* and time *t* is simulated from multinomial distribution with probability vector ***c***_*i*_(*t*) and a randomly simulated sequencing depth as the largest integer smaller than a *Unif* (5000, 50000) random variable. For the *i*th individual, longitudinal outcomes *Y*_*i*_(*t*), observable compositional features ***X***_*i*_(*t*) and the time vector *t*, (*t* = 0, …, *m* - 1) are used in model fitting.

We considered multiple scenarios which focused on different aspects of simulated data characteristics. For simulations with continuous outcomes, we set the reference scenario with *n* = 50, *p* = 200, *m* = 3, *u* = 0.15, *ρ* = 0 and *κ* = 2.5, which was based on the empirical observation of strong time effect on longitudinal outcome variables (such as BMI or dietary intake). Then we performed simulations with *n* = 10, 20, 50, 100, 200, *p* = 100, 200, 500, *m* = 2, 3, 4, 6, 8, *u* = 0.1, 0.15, 0.25, 0.5, *ρ* = 0, 0.4, 0.8 and *κ* = 0, 1.5, 2.5 while fixing other parameters as specified in the reference scenario. For simulations with binary outcomes, we set the reference scenario with *n* = 60, *p* = 200, *m* = 3, *u* = 0.3, *ρ* = 0 and *κ* = 0, where the sample size and effect size were higher than the reference scenario used for simulating the continuous outcome. This was because models for binary outcomes requires a larger sample size or effect size to reach similar level of power as models for continuous models. We let *κ* = 0 in the reference scenario for binary outcome simulations because a strong time effect would result in very small variations of a longitudinal binary variable (i.e. constantly equal to zero or one) as time increases. Centered at the reference scenario, we conducted simulations with *n* = 20, 30, 60, 100, 200, *p* = 100, 200, 500, *m* = 2, 3, 4, 6, 8, *u* = 0.15, 0.3, 0.45, 0.75, *ρ* = 0, 0.4, 0.8 and *κ* = 0, 1.5, 2.5.

#### 2.4.2 Real data examples

**Longitudinal diet and microbiome data of the NUTRIVENTION study** Elevated BMI and diets lacking plant foods are significant risk factors for multiple myeloma, which led to the development of a high fiber dietary intervention strategy. The MSK NUTRIVENTION study (NCT04920084) was a prospective trial investigating the efficacy of a high-fiber dietary intervention on weight loss and also whether it may delay progression from monoclonal gammopathy or smoldering myeloma to multiple myeloma [4]. The study recruited 20 evaluable patients who received 12 weeks of high fiber plant-based meals and 24 weeks of nutrition coaching with the meals and were followed for a year. Various patient characteristics, including but not limited to BMI, dietary intake, and 16S rRNA sequencing data from stool samples, were collected at 5 planned time points (baseline, 1 month, 3 months, 6 months, and 1 year) across a whole year of intervention. It has been shown that the intestinal alpha diversity was significantly increased from baseline to 3 months after study, where the longitudinal intestinal alpha diversity also had a significantly negative association with BMI [4]. The study showed that a high fiber dietary intervention could improve BMI and reshape the gut microbiome.

We applied the proposed method and other existing methods to correlate the longitudinal microbial taxa abundance with the longitudinal fiber intake collected from the 20 precursor plasma cell disorder patients receiving the high-fiber food intervention. 65 matched pairs of stool samples and fiber intake data points between baseline and 6 months after intervention were identified for the analysis, where each patient contributed 2-4 matched data points with a median of 3 per patient. The distribution of the available fiber intake values had a slightly heavy tail (**Fig.2A**), where the fiber intake was the highest 1 month after the start of intervention (**Fig.2C**). Such trajectories of fiber intake reflected patients’ adherence to the intervention protocol, where the adherence was the highest shortly after the study was initiated. Based on the above observations, we conducted natural logarithm transformation to the grams of fiber intake for modeling (**Fig.2B**), while using natural cubic spline terms to capture the non-linear time effect. **Fig.2D** displays the distribution of taxa prevalence, where taxa with prevalence below 10% are excluded. It can be seen that there are many taxa with high prevalence across samples.

**Figure 2:**
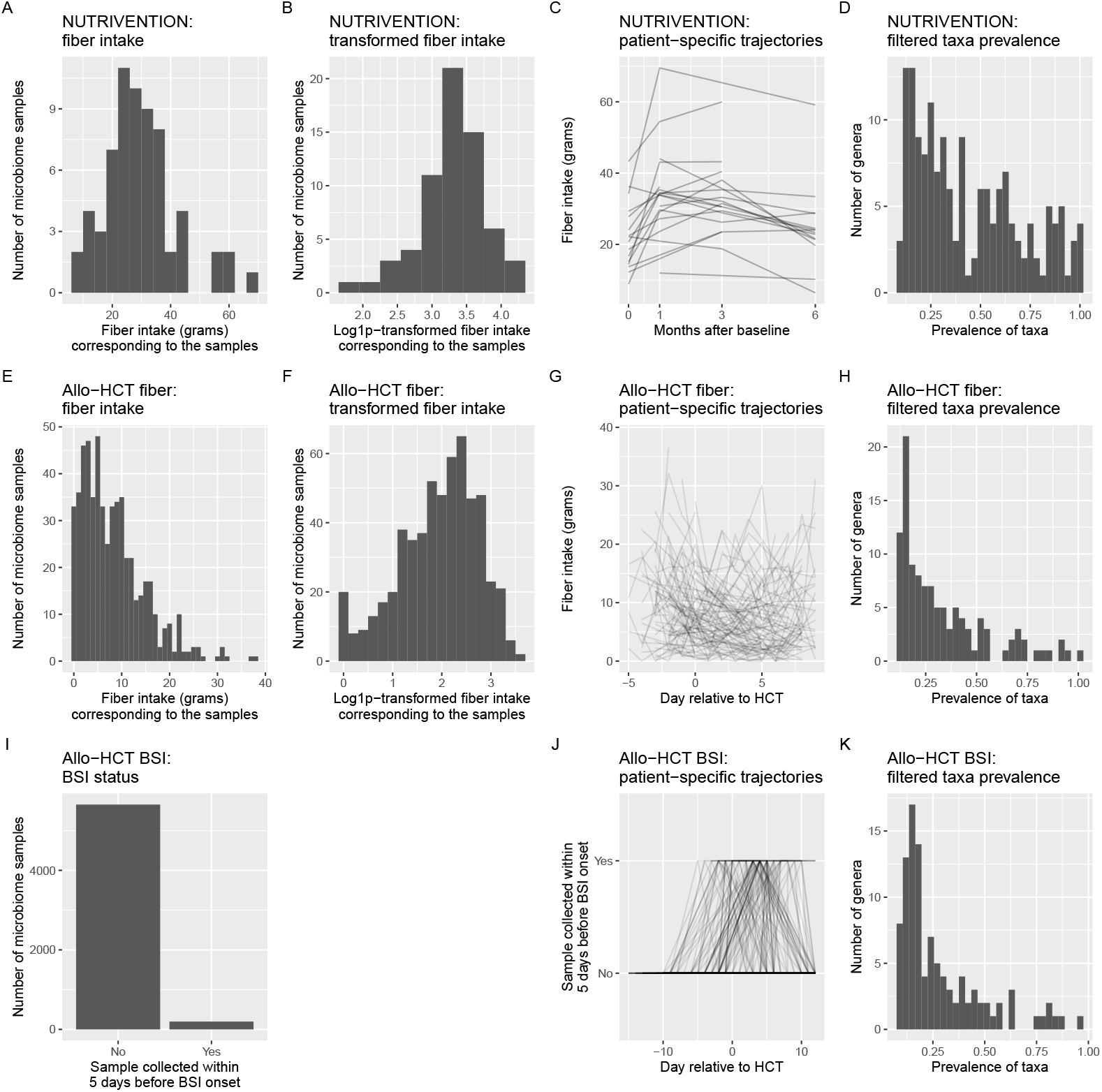
Data characteristics for the distribution of fiber intake values, fiber intake trajectories, and prevalence of filtered genera features. **A-D.** Histogram of fiber intake in grams (**A**), histogram of log-transformed fiber intake (**B**), spaghetti plot of patient specific fiber intake trajectories (**C**), and histogram of filtered taxa’s prevalences (**D**) for patients in the NUTRIVENTION study. **E-H**. Histogram of fiber intake in grams (**E**), histogram of log(·+1)-transformed fiber intake (**F**), spaghetti plot of patient specific fiber intake trajectories (**G**), and histogram of filtered taxa’s prevalences (**H**) for patients in the allo-HCT fiber cohort. **I-K**. Bar plot of sample-specific BSI status (**I**), spaghetti plot of BSI status based on available stool samples from different patients (**J**), and histogram of filtered taxa’s prevalences (**K**) for patients in the allo-HCT BSI cohort.

**Longitudinal diet and microbiome data of MSK allo-HCT cohort** To investigate the associations between dietary intake and the change of gut microbiota during bone marrow transplantation, the investigators at MSK collected longitudinal diet and 16S rRNA microbiome data for allo-HCT patients during the period of inpatient stay [10]. Specifically, food intake was categorized as five macronutrients (sugar, fiber, fat, protein, and other carbohydrates) in grams based on receipts from cafeteria and records from the care team to reflect food items and the amount of actual intake of each food item during each recorded meal. Accordingly, longitudinal stool samples were also collected for 16S rRNA sequencing. Based on a Procrustes analysis between microbial and macronutrient compositions, the alignment was the highest between between dietary records and stool samples collected 2 days later, as compared to other choices of gap days [10], which makes it possible to pair each dietary record with a stool sample based on the availability.

To demonstrate the utility of the proposed log-ratio PGEE method, we treated fiber intake in grams as the outcome variable in the GEE model, while using the genera counts observed in the corresponding stool sample collected two days later as covariates, aiming to identify microbial markers associated with fiber intake. The analysis focused on the diet records collected between 7 days prior to transplantation (day −7) and 7 days after transplantation (day 7), which consisted of 505 patient-days of paired fiber-microbiome longitudinal samples from 137 unique patients collected between day −5 and day 9. The number of available paired samples varied across patients, ranging from a single pair to 10 pairs with the median of 3 pairs and the interquartile range between 2 and 5 pairs. Due to the conditioning chemotherapy given before the HCT, overall dietary intake, including fiber, declined rapidly from day −7 to day 0 followed by a slow recovery trajectory after day 0 (**Fig.2G**). The heavy shift of dietary pattern also caused highly skewed distribution of fiber intake with a long tail (**Fig.2E**). Therefore, we conducted natural logarithm(+1) transformation to the fiber intake values (**Fig.2F**) in the models and adjusted for the non-linear time effect by including the cubic natural spline of time as covariates. Due to the conditioning therapy prior to the transplantation, patients’ gut microbiota were heavily shifted during the time window, where the prevalence was low for most taxa (**Fig.2H**).

**Longitudinal BSI and microbiome data of MSK allo-HCT cohort** Bloodstream infection (BSI) is a high-risk complication for allo-HCT patients, which has been shown to be related to the composition of gut microbiota [1]. For a cohort of 1,278 MSK allo-HCT patients, the longitudinal microbiome data and the corresponding BSI status information have been made publicly available [22]. Notably, more than 80% out of the 269 observed BSI events observed between day −14 and day 12 relative to transplantation were caused by *Enterococcus* species (55%) and *Escherichia coli* (28%).

To demonstrate FLORAL’s utility in handling a binary outcome variable in real data analysis, we focused on a subcohort of 1,206 patients with 5,851 available stool samples collected between day −14 and 12 relative to transplantation. Microbial features from the corresponding 16S sequencing data were correlated with the binary outcome variable of sample-specific BSI status, which was defined as whether or not collected within a 5-day window before the onset of BSI event. The number of samples per patient ranged between 1 and 25 with the median of 3 samples and the interquartile range between 2 and 6 samples. Due to the low prevalence of the BSI event, only 197 samples from 104 patients were collected within a 5-day window before the BSI event (**Fig.2I**), where the BSI events occurred between day −5 and day 12 relative to transplantation (**Fig.2J**). A similar distribution of taxa prevalence was observed between the allo-HCT fiber cohort (**Fig.2H**) and the allo-HCT BSI cohort (**Fig.2K**), showing consistencies of the study population of allo-HCT patients.

#### 2.4.3 Method configurations and assessment

**Simulations** We applied the proposed log-ratio PGEE model with two-step variable selection and other existing methods to benchmark the variable selection performance in simulations and real-data analysis. In simulation studies, log-ratio PGEE lasso models were fitted by the proposed method `FLORAL` with the correct compound symmetry working correlation structure (`FLORAL,cp`), and the incorrect independent (`FLORAL,ind`) and AR-1 (`FLORAL,ar1`) working correlation structures. In addition, we fitted standard PGEE models without the zero-sum constraint but with the correct compound symmetry working correlation structure using log-transformed count data (`PGEE,log`), relative abundance data (`PGEE,rel`), and centered log-ratio (CLR) transformed data (`PGEE,clr`). For the above penalized regression methods, a linear time effect was included in the GEE models without penalties (*r* = 0). Variable selection was performed by identifying the non-zero regression coefficients at *λ*_min_ and *λ*_1se_ based on a five-fold cross validation, where the fold split was set as identical for all methods within the same run. In addition, the implementation of the PGEE method without the zero-sum constraint was fulfilled by running the `FLORAL` function with *γ* = 0 to ensure identical cross-validation and variable selection procedures. As illustrated in the Introduction section, only a few existing methods can be applied to study associations between longitudinal microbial features and longitudinal outcomes [13–15]. We implemented the mixed-effect model `MaAsLin2` [13] and `MaAsLin3` [14] with CLR transformation (`normalization=‘CLR’, transform=‘NONE’`) or with total sum scaling (TSS) normalization and log-transformation (`normalization=‘TSS’,transform=‘LOG’`). For both methods, the mixed-effect models considered a linear time effect, a linear outcome effect from the simulated longitudinal outcome, and a random intercept for each simulated individual. Feature selection was based on FDR-adjusted p-values via the Benjamini-Hochberg approach [32] with significance level of 0.1. For `MaAsLin3`, we report features selected by the prevalence model and the abundance model separately.

In terms of method assessment, 100 runs were performed under each simulation scenario for performance evaluations. Four commonly applied metrics were calculated for each method, namely the *F*_1_ score, number of false positive features, number of false negative features, and false discovery rate (FDR). The *F*_1_ score is defined as the harmonic mean of precision (positive predictive value) and recall (sensitivity), which is used to indicate the overall variable selection performance after balancing sensitivity and FDR. An *F*_1_ score of 1 indicates perfect performance, while an *F*_1_ score of 0 implies that no true features were selected. The cross-validated deviance residuals were also compared across the `FLORAL` models with different working correlation structures to evaluate its effectiveness of selecting models with the most appropriate working correlation structure. In addition, we also recorded the time (in seconds) used for each evaluated method to complete each simulation run, the number of iterations `FLORAL` took before convergence or reaching the maximum number of iterations, and the convergence criterion 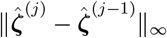 when `FLORAL’s` algorithm (**Algorithm 1**) stopped at each simulation run.

**Real-data analysis** As discussed in Section 2.4.2, we performed natural log-transformation to normalize the fiber intake data in the NUTRIVENTION and allo-HCT fiber cohort, where the transformed fiber intake was treated as a continuous outcome variable. For the allo-HCT BSI cohort, the BSI status was used as a binary outcome variable. We fitted the proposed log-ratio PGEE models with independent (`FLORAL,ind`), compound symmetry (`FLORAL,cs`), and AR-1 (`FLORAL,AR1`) working correlation structures. In addition, we also applied PGEE without zero-sum constraints with log-transformed (`PGEE,log`), relative abundance (`PGEE,rel`), and CLR-transformed (`PGEE,clr`) microbiome data. Cubic natural spline terms were included for all three datasets as covariates without penalization (*r* = 0), where the knots were selected as the 10th, 50th and 90th quantiles of the time points of sample collection. Variable selection procedure follows the same procedure as described for the simulation studies with a 5-fold cross validation. Additionally, we conducted model fitting using the above PGEE methods for 100 times with random fold splits, then summarized the number of times for taxa being selected across 100 times as probabilities of being selected. Taxa with high frequency of selection will be interpreted as more likely to be associated with the fiber intake or BSI onset. We also applied mixedeffect models `MaAsLin2` and `MaAsLin3` by treating transformed fiber intake or binary BSI status as a covariate. Similar to the penalized regression models, we also included the same cubic natural spline terms to capture non-linear time effects. For the MaAsLin packages, we applied the two taxa normalization-transformation configurations as used in simulations. Selected features are defined as features with FDR-adjusted p-values *<* 0.1. We also report both prevalence and abundance models for MaAsLin3.

We evaluated the methods based on their capabilities of detecting signals and the clinical relevance of selected microbial features. We also compared the features with the strongest signals from different methods, as ranked by the selection probabilities for the PGEE models and the p-values for the mixed-effect models.

## 3 Results

### 3.1 FLORAL achieves superior variable selection performances in simulations

We performed extensive simulations to assess the variable selection performance of the proposed log-ratio PGEE method `FLORAL`, the standard `PGEE` models with log-transformed, relative abundance, and CLR-transformed features, and mixed-effect models `MaAsLin2` and `MaAsLin3`. For both continuous and binary outcome simulations, we specified the compound symmetry (`cs`) working correlation structure for data generation. In model fitting, we tested using the correct structure (`cs`), the independence structure (`ind`) and the AR-1 structure (`ar1`) with `FLORAL`, while the model fitting with other PGEE methods were conducted with the correct correlation structure (`cs`). Details of data generation and performance assessment can be found in Section 2.4. **Fig.3** summarizes the median *F*_1_ scores obtained by the PGEEs and mixed-effect models across 100 simulations for each scenario. Overall, the task of variable selection is more challenging for longitudinal binary outcomes as compared to longitudinal continuous outcomes, such that the simulations with continuous outcomes attained better performance than those with binary outcomes even with smaller sample sizes or effect sizes (Figs.3A-C). The above observation justifies our simulation strategies with different reference scenario for continuous and binary outcomes.

**Figure 3:**
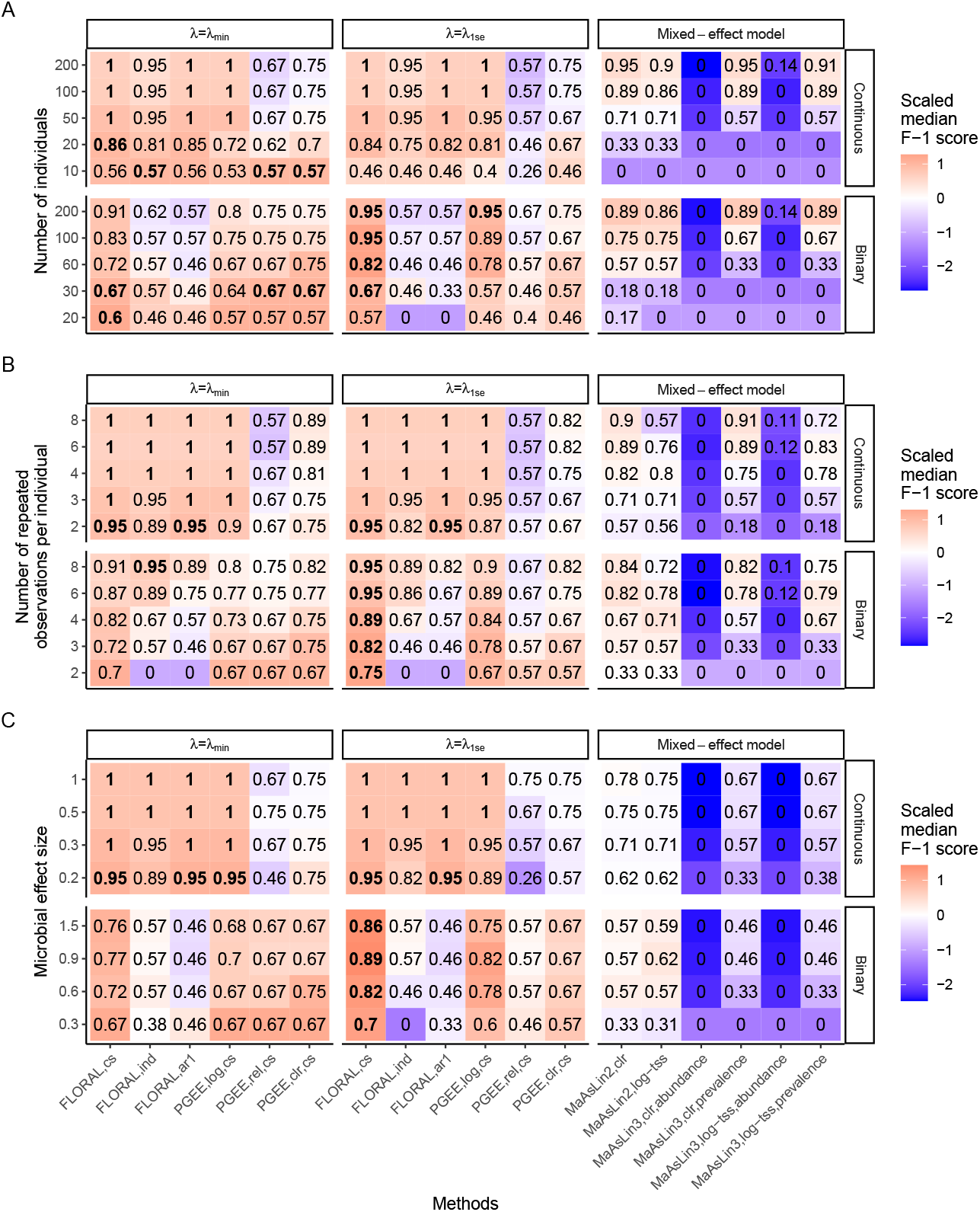
Median *F*_1_ scores out of 100 simulations obtained by PGEE methods with *λ* = *λ*_min_, *λ* = *λ*_1se_, and mixed-effect models (MaAsLin) under different simulation scenarios with continuous or binary longitudinal outcome variables with different **A.** number of individuals (*n*), **B**. number of repeated observations per individual (*m*) and **C**. microbial feature effect sizes (*u*). As described in the Method section, the reference scenario is *n* = 50, *p* = 200, *m* = 3, *u* = 0.15, *ρ* = 0, *κ* = 2.5 for continuous outcome and *n* = 60, *p* = 200, *m* = 3, *u* = 0.3, *ρ* = 0, *κ* = 0 for binary outcome. For each scenario, the color scheme represents scaled median *F*_1_ scores, where red color represents better performance. The highest *F*_1_ score per scenario was shown in bold fonts. For the PGEE methods, we considered compound sysmmetry (cs), independence (ind) and AR-1 (ar1) working correlation structures.

Comparing across the methods, most PGEE methods conducted better variable selection than the mixed-effect models under small numbers of individuals (*n*), repeated observations (*m*), and effect sizes (*u*), while the mixed-effect models achieved comparable performances as *n* ≥ 100 (**Fig.3**,**Figs.S1-S3**, panel A). Specifically, the PGEE methods showed higher sensitivity in selecting the true features than the mixed-effect models under smaller sample sizes and effect sizes (**Figs.S1-S3**, panels C). Moreover, the FDR and false-positive control of the PGEE methods gradually improved with a higher sample size, while we observed an inflated FDR of `MaAsLin2` and `MaAsLin3` as *n* and *m* increased (**Figs.S1-S2**, panels D). Additional simulations also implied that an increasing number of features (*p*) corresponded to a decline in sensitivity or an inflation in FDR for all methods, resulting in a decreasing *F*_1_ score (**Fig.S4**). In addition, feature-wise correlation level (*ρ*) and the strength of the linear time effect (*κ*) appeared not to heavily affect the variable selection performance (**Figs.S5-S6**).

Among the PGEE methods applying the underlying correct compound symmetry working correlation structure, `FLORAL` achieved a consistently better balance between sensitivity and FDR control while keep both in reasonably effective levels, resulting in a better overall *F*_1_ score. Due to the log-ratio model used for data generation, `FLORAL` and the standard PGEE model with log-transformed data (`PGEE-log`) achieved comparably high level of sensitivity than the PGEE models using other data transformation schemes. Nevertheless, the `FLORAL` model obtained slightly higher sensitivity and consistently better FDR control than `PGEE-log` in most scenarios (**Figs.S1-S6**, panels C-D), where the improved sensitivity can be attributed to the implementation of zero-sum constraint to account for compositionality, and the better FDR control is due to the additional two-step feature screen strategy. The performances of `FLORAL` with *λ*_min_ and *λ*_1se_ penalty parameters were generally similar, where *λ*_min_ tended to select more truly associated features at the cost of inflated false positive findings, while *λ*_1se_ tended to be more conservative with a well-controlled FDR. Overall, `FLORAL` achieved effective FDR control for continuous outcomes when the sample size satisfies *n* ≥ 20, irrespective of the choice of *λ*. However, we did observe a more severely inflated FDR associated with *λ*_min_ for binary outcomes in most simulation scenarios, whereas `FLORAL` with *λ*_1se_ still maintained a decent level of FDR with binary outcomes (**Figs.S1-S6**, panel D).

`FLORAL` achieved robust variable selection performance when using different working correlation structures in modeling continuous outcomes, while the performance for binary outcomes depended more heavily on the specification of correct working correlation structure. As shown in **Fig.3** and **Figs.S1-S6** panel A, the performance of variable selection in the continuous outcome models was not strongly affected by the specification of working correlation, where `FLORAL,cs` and `FLORAL,ar1` reached slightly higher *F*_1_ scores than `FLORAL,ind` due to better sensitivity (**Figs.S1-S6**, panel C). In simulations with binary outcomes, on the other hand, we observed a large performance gap between PGEE models with correctly specified working correlation and `FLORAL` with incorrectly specified correlation structures. Although the overall performance of `FLORAL,ind` and `FLORAL,ar1` improved with larger sample sizes, the sensitivity of variable selection was consistently lower than other PGEE methods with correctly specified working correlation structure (**Figs.S1-S6**, panel C). The above observations align with the comparisons of cross-validated deviance residuals across the three working correlation structures, where all three structures achieved comparable cross-validated deviance residuals with continuous outcome, while the compound symmetry structure obtained much smaller deviance residuals than the other two structures (**Fig.S7-S12**, panel B). Such alignment demonstrated the utility of cross-validated deviance residuals for model selection.

The two mixed-effect models, `MaAsLin2` and `MaAsLin3`, required a larger sample size to achieve comparable performances as compared to `FLORAL`. Due to the data generation mechanism based on a log-ratio model, mixed-effect models with CLR-transformation (clr) performed better than models with log-transformed TSS (`log-tss`) in most scenarios (**Figs.S1-S6**, panel A). In addition, the high sparsity with 80% zeros of simulated microbial features resulted in more informative variable selection results from `MaAsLin3`’s prevalence model as compared to the abundance model (**Fig.3**). Moreover, `MaAsLin2` and both models from `MaAsLin3` tended to have inflated FDRs under simulations with large *n* and large *m* (**Figs.S1-S2**, panels B,D), which was also described by the preprint of MaAsLin3 as “precision loss with high power” [14, Figs.S4 and S7].

In terms of computational time, `FLORAL` generally took a longer time than `MaAsLin2` but a shorter time than `MaAsLin3` (**Figs.S7-S12**, panel A). As observed, FLORAL requires more computational time under simulations with larger number of features (**Fig.S10A**). Moreover, `FLORAL` typically took less than 50 iterations to converge under *λ* = *λ*_min_ and *λ* = *λ*_1se_, where models for binary outcomes may take more iterations than models for continuous outcomes (**Figs.S7-S12**, panel C). In very rare cases, `FLORAL` did not reach convergence after 100 iterations, while the convergence criterion was not distant from the pre-specified threshold of 0.001 (**Figs.S7-S12**, panel D).

### 3.2 FLORAL identifies meaningful taxonomic markers associated with the fiber intake of cancer patients

We correlated longitudinal fiber intake records and longitudinal microbial genera from two cancer studies. The NUTRIVENTION study [4] is a pilot trial with 20 patients and less frequent sample collections, while the MSK allo-HCT cohort [10] is a larger cohort with more than 100 patients and more frequent sample collections. To benchmark the feature selection performance of various methods under different data characteristics, we identified matched fiber intake and microbiome data points for the NUTRIVENTION study (65 samples from 20 patients with 161 genera) and the MSK allo-HCT cohort (505 samples from 137 patients with 112 genera), where the included genera were detected in more than 10% of all samples. Similar to the simulations, we applied `FLORAL` with three working correlation structures (`cs, ind` and `ar1`), the standard PGEE models with cs correlation structure with log-transformed, relative abundance, and CLR-transformed data, and mixed-effect models (`MaAsLin2` and `MaAsLin3`). The `FLORAL` and PGEE models were run for 100 times with random fold split to reflect a robust pattern of feature selection by the 5-fold cross-validation, where the more frequently selected taxa indicate stronger signals. We used the threshold of 0.1 of the adjusted p-values for feature selection from the mixed-effect models. Detailed information about the two studies and method configurations can be found in Section 2.4.

### FLORAL identifies gut health indicating genera from NUTRIVENTION data

**Fig.4** displays the variable selection results of `FLORAL` for the NUTRIVENTION data, where features were only identified using *λ* = *λ*_min_ due to the small sample size and limited statistical power. Cross-validated deviance residual implies similar model fitting performances by the three working correlation structures (**Fig.4A**), which is confirmed by the correspondingly similar variable selection results (**Fig.4B-D**). Genera *Coprococcus* and *Longicatena* were selected in more than 70% of cross-validated runs, where *Coprococcus* is a well-studied butyrate producer that secretes beneficial short chain fatty acids [33] and *Longicatena* is associated with gut dysbiosis and inflammatory bowel disease [34]. As expected, the abundance of *Coprococcus* was positively associated with fiber intake while the abundance of *Longicatena* was negatively associated with fiber intake, indicating the fiber-oriented dietary intervention was effective in boosting beneficial bacteria and controlling potential pathogens. Genera *Longibaculum* was also identified as positively associated with fiber intake in around 40% of runs, which has been shown to improve oral glucose tolerance in a mouse study [35].

**Figure 4:**
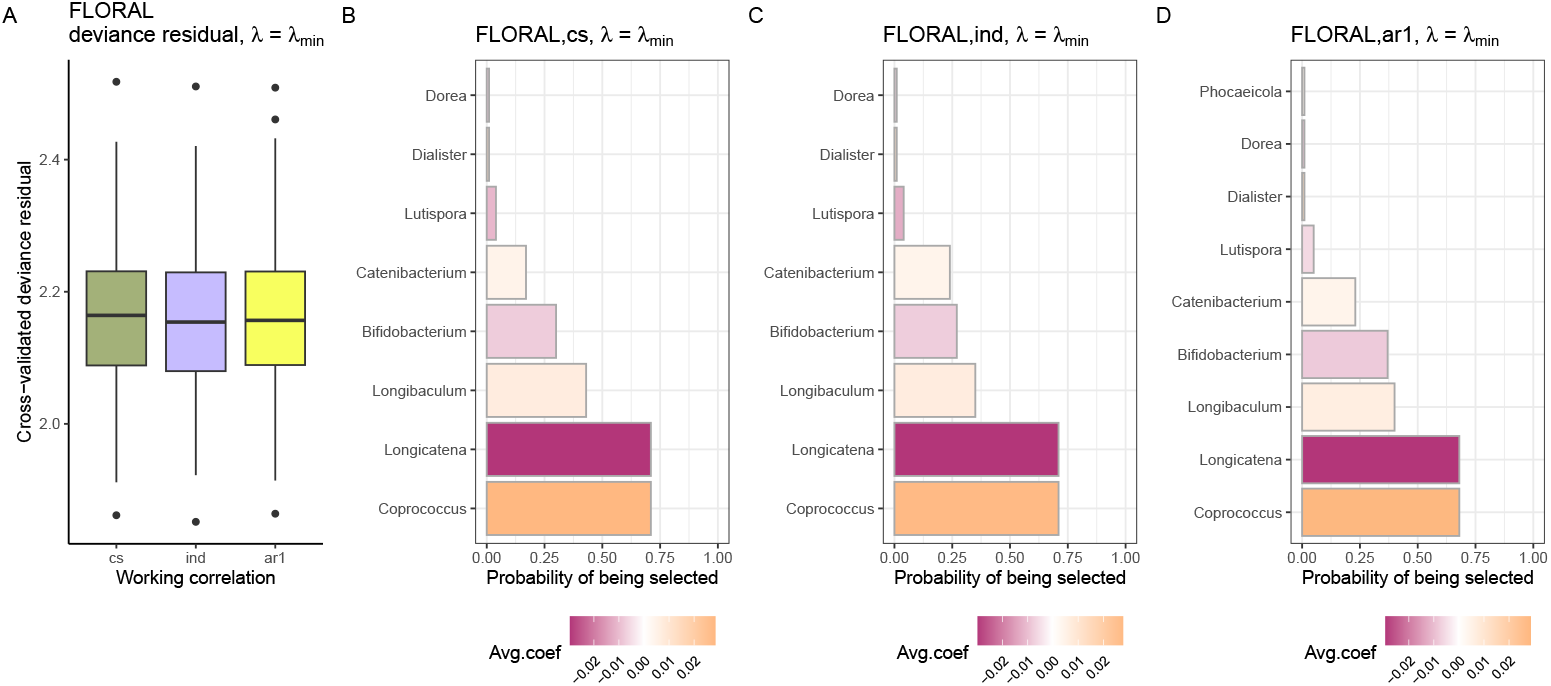
Model fitting and variable selection results for the NUTRIVENTION longitudinal fiber intake and microbiome data using FLORAL with *λ* = *λ*_min_. **A.** Cross-validated deviance residual obtained by `FLORAL` using compound symmetry (cs), independence (ind) and AR-1 (ar1) correlation structures out of 100 runs of 5-fold cross-validation with random fold splits. **B-D**. Proportions of taxa being selected by `FLORAL` out of 100 runs of 5-fold cross-validation with random fold splits using **B**. compound symmetry, **C**. indpendence and **D**. AR-1 working correlation structures. Colors represent the average feature coefficient out of 100 runs, where a positive coefficient implies a positive association between fiber intake and the microbial feature. Results with *λ* = *λ*_1se_ were omitted as no features were selected.

Out of the standard PGEE models with different data transformation schemes, only the model using log-transformed count data identified several markers with *λ* = *λ*_min_ (**Fig.S13A**). This observation is consistent with our simulation studies where PGEE models with relative abundance and CLR-transformed data showed poorer sensitivities compared to `FLORAL` and PGEE with log-transformed count data (**Fig.S1-S6**, panel C). Similar to `FLORAL`, PGEE with log-transformed data also selected *Coprococcus* and *Longicatena*. However, the PGEE model did not adjust for varying sequencing depths across samples for the log-transformed taxa counts due to the lack of zero-sum constraint, which further caused the under-selection of *Coprococcus* and over-selection of *Longicatena*. In terms of the mixed-effect models, only `MaAsLin2` with CLR-transformed data selected four taxa associated with fiber intake at the FDR threshold of 0.1 (**Fig.S13B**), where fiber intake was positively associated with *Anaerofilum* (q=0.07) and *Coprococcus* (q=0.08) and negatively associated with *Longicatena* (q=0.09) and *Dehalobacter* (q=0.10). While the clinical interpretation for Coprococcus and Longicatena is expected, the clinical interpretation for Anaerofilum is unclear given there is evidence for its association with obesity [36].

### FLORAL identifies clinically relevant genera from MSK allo-HCT data

**Fig.5** shows model fitting and variable selection results by `FLORAL` for the MSK allo-HCT fiber cohort. Compared to the NUTRIVENTION data, the allo-HCT fiber cohort consists of substantially more samples and patients, where features were selected using both *λ* = *λ*_min_ and *λ* = *λ*_1se_ configurations. Similar to the simulations for continuous outcomes and the NUTRIVENTION analysis, different working correlation structures again reached a similar level of cross-validated deviance residual (**Fig.5A,E**), showing similar model fitting and variable selection performances (**Fig.5B-D,F-H**). Combining the results from *λ* = *λ*_min_ and *λ* = *λ*_1se_ across different working correlation structures, an increasing fiber intake was most strongly associated with an increasing abundance of *Blautia* and decreasing abundances of *Enterococcus, Alistipes* and *Veillonella*. Out of the above four markers with strongest associations, *Blautia* and *Enterococcus* have been extensively studied in allo-HCT literature as taxa associated with good and poor clinical outcomes, respectively [2, 37, 38]. Moreover, *Veillonella* is an oral bacteria and an indicator of gut microbiota depletion in allo-HCT patients [39], while *Alistipes* has been shown to have both protective and harmful effects on gut health [40], also offering reasonable interpretations of their associations with fiber intake.

**Figure 5:**
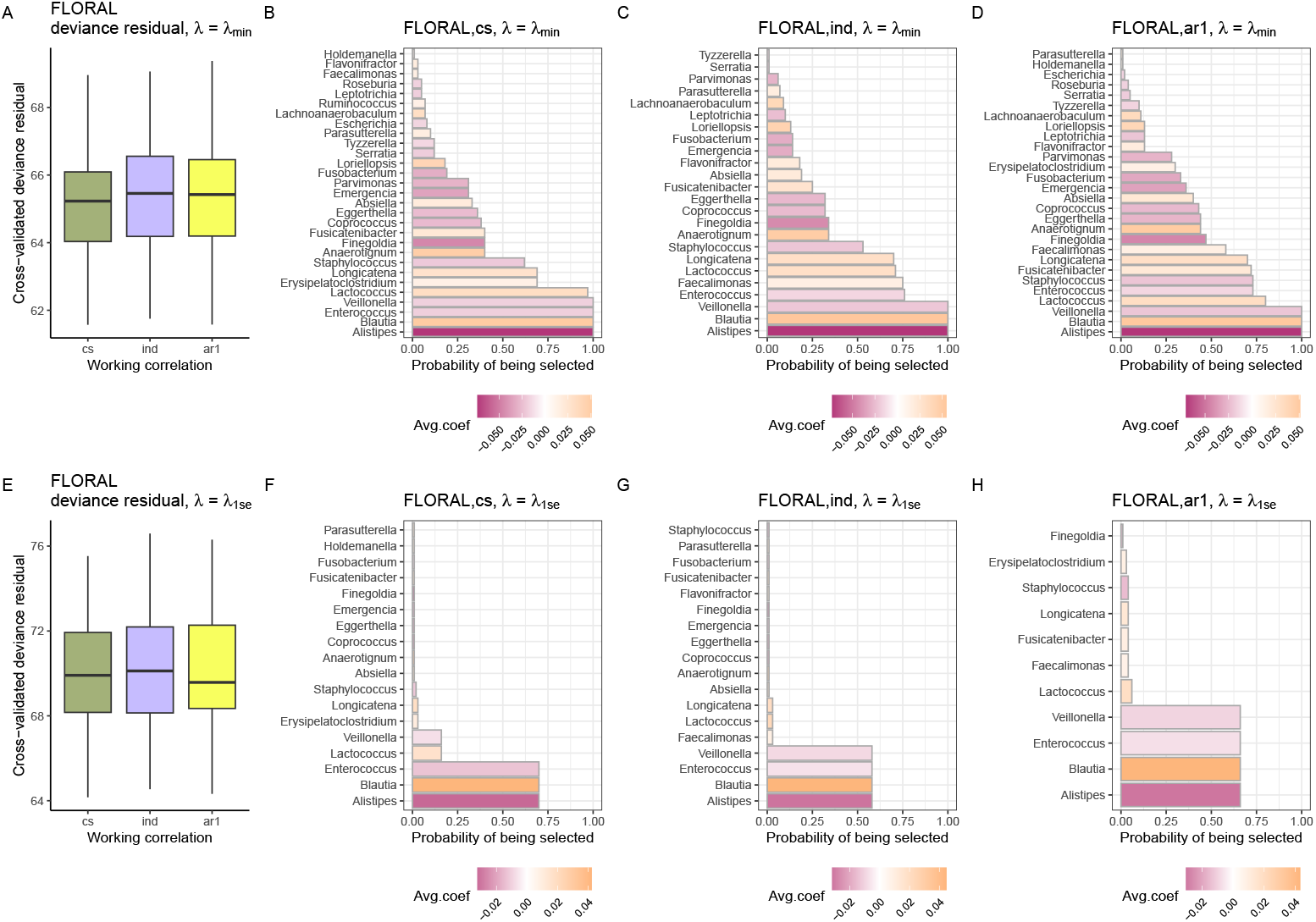
Model fitting and variable selection results for the allo-HCT longitudinal fiber intake and microbiome data using FLORAL with *λ* = *λ*_min_ (**A-D**) and *λ* = *λ*_1se_ (**E-H**). **A**,**E.** Cross-validated deviance residual obtained by `FLORAL` using compound symmetry (cs), independence (ind) and AR-1 (ar1) correlation structures out of 100 runs of 5-fold cross-validation with random fold splits. **B-D**,**F-H**. Proportions of taxa being selected by `FLORAL` out of 100 runs of 5-fold cross-validation with random fold splits using **B**,**F**. compound symmetry, **C**,**G**. indpendence and **D**,**H**. AR-1 working correlation structures. Colors represent the average feature coefficient out of 100 runs, where a positive coefficient implies a positive association between fiber intake and the microbial feature.

Similar to the NUTRIVENTION analysis results, the PGEE model with log-transformed count data identified similar features as selected by `FLORAL` with different ranks of selection frequencies (**Figs.S14A**,**D**). Additionally, PGEEs with relative abundance and CLR-transformation also identified *Enterococcus* or *Blautia* in most runs with *λ* = *λ*_min_ (**Figs.S14B-C**), while still showing low sensitivities at *λ* = *λ*_1se_ (**Figs.S14E-F**). In terms of the mixed-effect models, both `MaAsLin2` and `MaAsLin3` identified multiple genera significantly associated with fiber intake at the FDR level of 0.1 (**Figs.S15-S16**), where feature selection was mainly driven by the models instead of the data transformation and normalization strategies. Interestingly, *Enterococcus* was not identified by either of the mixed-effect models, while *Blautia* was only identified by `MaAsLin2`, with the 3rd strongest association (q=0.015) per the `MaAsLin2,clr` model (**Fig.S16D**) and only the 14th strongest association (q=0.09) in the `MaAsLin2,log-tss` model (**Fig.S16A)**. While obtaining numerous significant associations, most of them had no established relevance with allo-HCT patient outcomes, making it challenging for generating new hypotheses.

### FLORAL precisely identifies BSI-causing taxa from MSK allo-HCT data

Model fitting and variable selection results by `FLORAL` for the MSK allo-HCT BSI cohort are shown in **Fig.6**. The independence working correlation structure obtained slightly lower cross-validated deviance residual compared to the other alternatives, while the variable selection results were highly similar across different model configurations. The two genera contributing to more than 80% of BSI events, *Enterococcus* and *Escherichia-Shigella*, were robustly selected as enriched in samples collected within 5 days before BSI onset by all models, irrespective of fold splits and the choice of penalty parameters. The above finding reveals a strong association between the composition of intestinal micro-biota and BSI. In addition, genera with health benefits, such as *Blautia* and *Lactococcus* were identified to be enriched in samples with BSI-negative status.

**Figure 6:**
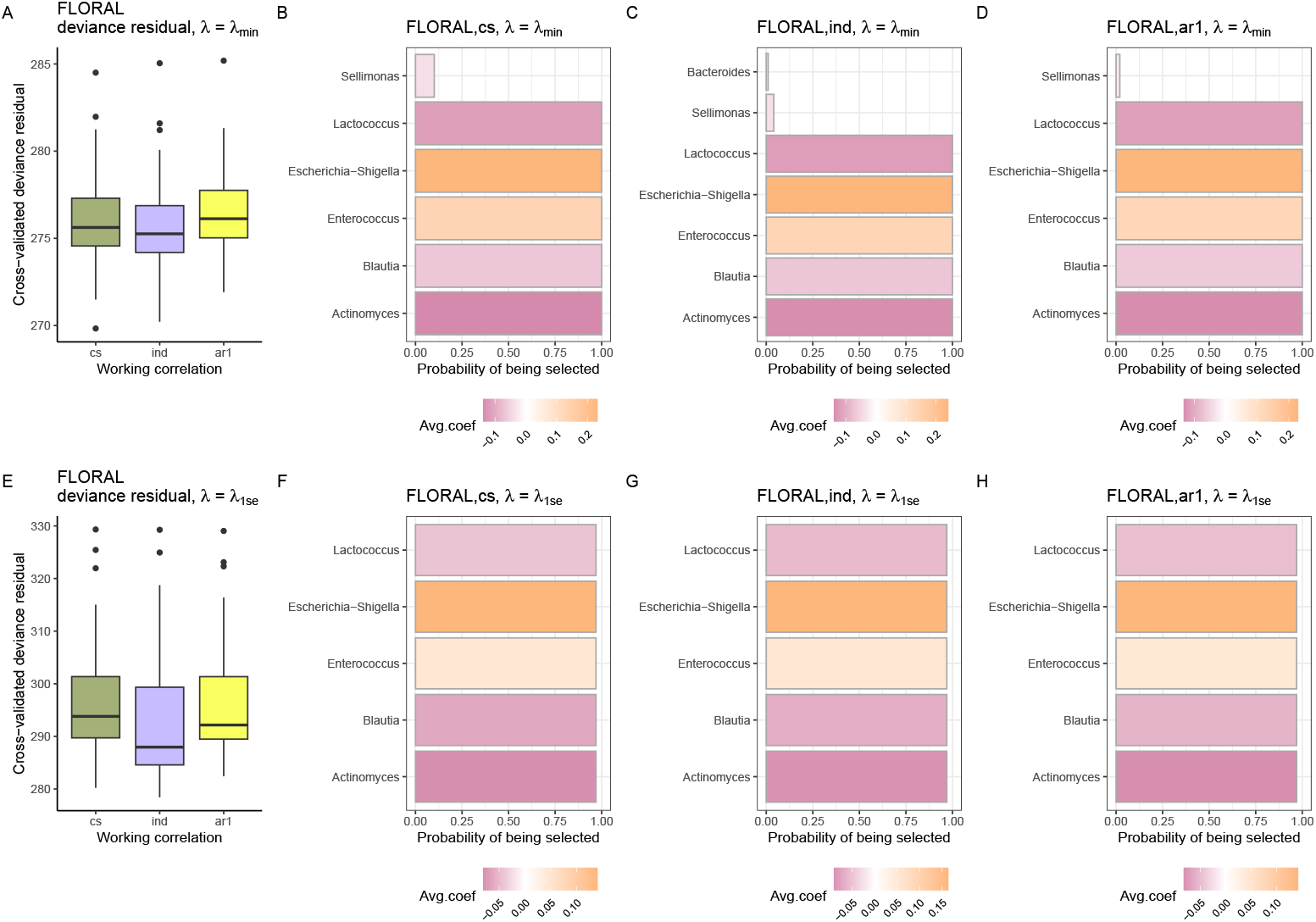
Model fitting and variable selection results for the allo-HCT longitudinal BSI status and microbiome data using FLORAL with *λ* = *λ*_min_ (**A-D**) and *λ* = *λ*_1se_ (**E-H**). **A**,**E.** Cross-validated deviance residual obtained by `FLORAL` using compound symmetry (cs), independence (ind) and AR-1 (ar1) correlation structures out of 100 runs of 5-fold cross-validation with random fold splits. **B-D**,**F-H**. Proportions of taxa being selected by FLORAL out of 100 runs of 5-fold cross-validation with random fold splits using **B**,**F**. compound symmetry, **C**,**G**. indpendence and **D**,**H**. AR-1 working correlation structures. Colors represent the average feature coefficient out of 100 runs, where a positive coefficient implies a positive association between a positive BSI status and the microbial feature.

PGEE models with log-transformed or CLR-transformed data also identified *Enterococcus* and *Echerichia-Shigella* as enriched in BSI-approching samples (**Fig.S17**). Features selected by the PGEE model with log-transformed data are highly similar to the selected features by `FLORAL` models (**Figs.S17A**,**C**), implying that the log-transformation may work without the zero-sum constraint when the signals are sparse and strong. In contrast, the PGEE model with CLR-transformed data selected additional taxa under *λ* = *λ*_min_ which might be false positives and did not reach as high selection frequency as FLORAL achieved under *λ* = *λ*_min_ and *λ*_1se_ (**Figs.S17B**,**D**). Moreover, models with relative abundance did not identify any features. Compared to the PGEE models, mixed-effect models identified a larger number of associations at the FDR level of 0.1 **Figs.S18-S19**, where the majority of identified features were enriched in BSI-negative samples. *Echerichia-Shigella* was identified by all `MaAsLin2` and `MaAsLin3` models as the most enriched taxa in BSI approaching samples (**Fig.S19**), while *Enterococcus* was identified as the second most significantly enriched taxon by `MaAsLin3,log-tss`’s abundance model (q=0.0001), the third most significantly enriched taxon by `MaAsLin2,log-tss` (q=0.01) and `MaAsLin3,clr`’s abundance model (q=0.003), the sixth most significantly enriched taxon by `MaAsLin2,clr` (q=0.004), and was not selected by the prevalence models of MaAsLin3 (**Figs.S19B**,**E**). Among models which did not rank *Enterococcus* as the second most significantly enriched taxon in BSI approaching samples, genera including *Fusobacterium, Phascolarctobacterium, Leptotrichia*, and *Finegoldia* were ranked ahead of *Enterococcus* but were not recorded as an infectious agent of actually observed BSI events. Comparing across the tested methods in real-data analyses, `FLORAL` achieved desirable feature-selection performance not only by successfully identifying easily interpretable taxonomic markers, but also by effectively ranking the most plausible and relevant associations as the strongest signals. This can be an advantage in exploratory analysis with an aim of hypothesis generation, where the mixed-effect methods rely on subjective p-value thresholds and may not identify the most biologically meaningful markers as having the most significant associations.

## 4 Discussion

In this work we introduced the log-ratio penalized generalized estimating equation (PGEE) method to our recently described `FLORAL` package as a new approach to analyzing longitudinal associations between microbial features and patient characteristics. We account for the compositionality of microbial features by an added zero-sum constraint [23] to the standard penalized estimating equation framework [17, 18] and impose a two-step variable selection procedure to better control the FDR [20]. Our simulations demonstrated superior sensitivity with reasonable FDR control of the proposed method over standard PGEE methods and mixed-effect methods under our model assumptions. Real-data analyses further validated the utility of our method in reliably identifying clinically relevant and reported gut health indicating taxonomic markers. Unlike the mixed-effect models where the selected features may contain biologically irrelevant microbes, `FLORAL`’s log-ratio PGEE more robustly ordered the highly relevant taxa as the strongest signals, showing high potentials in exploratory analysis for longitudinal microbiome studies of various scales.

The proposed log-ratio PGEE method is naturally implemented in the publicly available R package `FLORAL`, which was first introduced for penalized log-ratio generalized linear models and Cox proportional hazards regressions [11]. We provide a user-friendly interface with a standard format of visualizations of variable selection results. Our simulations also validated the stability of the proposed minorize-maximize algorithm with a zero-sum constraint by showing high convergence rates and fast computational speed. Moreover, we propose a built-in model selection criterion for different working correlation structures based on cross-validated deviance residual, which performs robustly in simulations.

Unlike the popularly applied mixed-effect models, we treat the longitudinal patient characteristic of interest as the outcome variable in a GEE model, which translated the research question into fitting a single model rather than hundreds of taxon-specific models. We believe this strategy is technically simpler and performance-wise more robust in real-data applications than the taxon-specific modeling approach, as the microbial trajectories are usually highly heterogeneous and are challenging to be explained by a unified set of model configurations. Under our setting, an analyst has the bandwidth to focus more carefully on modeling the single trajectory of the “outcome” variable of interest, such as dietary intake, which usually follows a more regular distribution than the sparse and skewed microbial abundance, and is easier for fine tuning the non-linear associations with respect to the time effect. Additionally, modeling an individual taxon usually involves multiple factors such as dietary intake, antibiotics, and other medications, where the colinearity between factors can easily mask the signals. Using FLORAL’s unified GEE model, on the other hand, we are positioned to concentrate more on the associations between the outcome variable of interest and microbial features, where the confounding factors to the trajectory of the outcome variable can still be adjusted. Furthermore, the proposed PGEE method can also be applied to model the trajectory of the outcome variable conditioned on a single baseline microbiome sample, which is another widely available data structure, especially in mouse studies.

Like other penalized regression methods, the proposed log-ratio PGEE model has the following limitations. First, the proposed method tends to have an inflated FDR with *λ* = *λ*_min_ when the number of individuals is small (**Fig.S1D**) and the number of features is large (**Fig.S4D**) especially for binary outcomes. If strict FDR control is desired, we recommend using *λ* = *λ*_1se_ for better FDR control at the cost of getting lower sensitivity. More systematic FDR control procedures, like knockoff [41], can be considered as a direction for future development. Second, the log-ratio regression framework cannot be easily extended to account for non-linear associations between microbial features and the outcome variable, which could be better captured by quantile-based methods [15] or dimension reduction methods for microbial trajectories [12]. Third, our implementation of the log-ratio PGEE model does not incorporate statistical inference of the regression coefficients as discussed in the original PGEE paper [18], where the construction of variance estimators and inference procedures can be further studied for log-ratio-based regression models.

## Data and Code Availability

Open-source R package `FLORAL` can be accessed via GitHub (https://vdblab.github.io/FLORAL) or CRAN (https://cran.r-project.org/package=FLORAL). Unpublished 16S rRNA sequencing data sets for NUTRIVENTION and MSK allo-HCT fiber will be made available on FigShare by the time of publication. 16S rRNA sequencing dataset for the MSK allo-HCT BSI cohort can be downloaded from https://doi.org/10.6084/m9.figshare.13584986 [22].

## Author Contributions

Teng.F. conceived of the project, developed the methodology and wrote the manuscript. Teng.F. and V.D. performed computational analysis. Teng.F., V.D., Tyler.F., M.B., N.R.W., A.D., S.S.R, U.A.S. and J.U.P analyzed and interpreted the analysis results. Tyler.F., M.B., N.R.W., J.P., A.D., F.C., J.H., A.G, S.S.R., U.A.S. assisted with microbiome and clinical data harmonization. J.P., F.C., J.H., A.G., S.S.R, A.M.L., U.A.S., M.R.M.v.d.B, and J.U.P. coordinated clinical data and sample collection and sequencing management. Teng.F., U.A.S., M.R.M.v.d.B. and J.U.P. co-supervised the study.

## Acknowledgments

This research was supported by National Cancer Institute award numbers P30-CA008748 MSK Cancer Center Support Grant/Core Grant, R25-CA272282, and R01-CA228308. Additional support was received from National Cancer Institute award numbers R01-CA228358, K12-CA184746 and R25-CA203650, National Heart, Lung, and Blood Institute (NHLBI) award numbers R01-HL123340, R01-HL147584 and K08-HL143189, National Institute on Aging award number P01-AG052359, The Allen Foundation, Inc, American Society of Hematology Scholar Award, Cycle for Survival, The Lymphoma Foundation, Parker Institute for Cancer Immunotherapy, Paula and Rodger Riney Multiple Myeloma Research Initiative, Seres Therapeutics, The Solomon Microbiome Nutrition and Cancer Program, Starr Cancer Consortium, The Susan and Peter Solomon Family Fund, Tri-Institutional Stem Cell Initiative, and V Foundation. The NUTRIVENTION trial was supported by Plantable that provided subsidized meals and participant nutrition coaching. We thank the MSK Society for the support of the Genomics Experience for Master’s Students (GEMS) program.

