## Supplemental Figures for "Correlating High-dimensional longitudinal microbial features with time-varying outcomes with FLORAL"

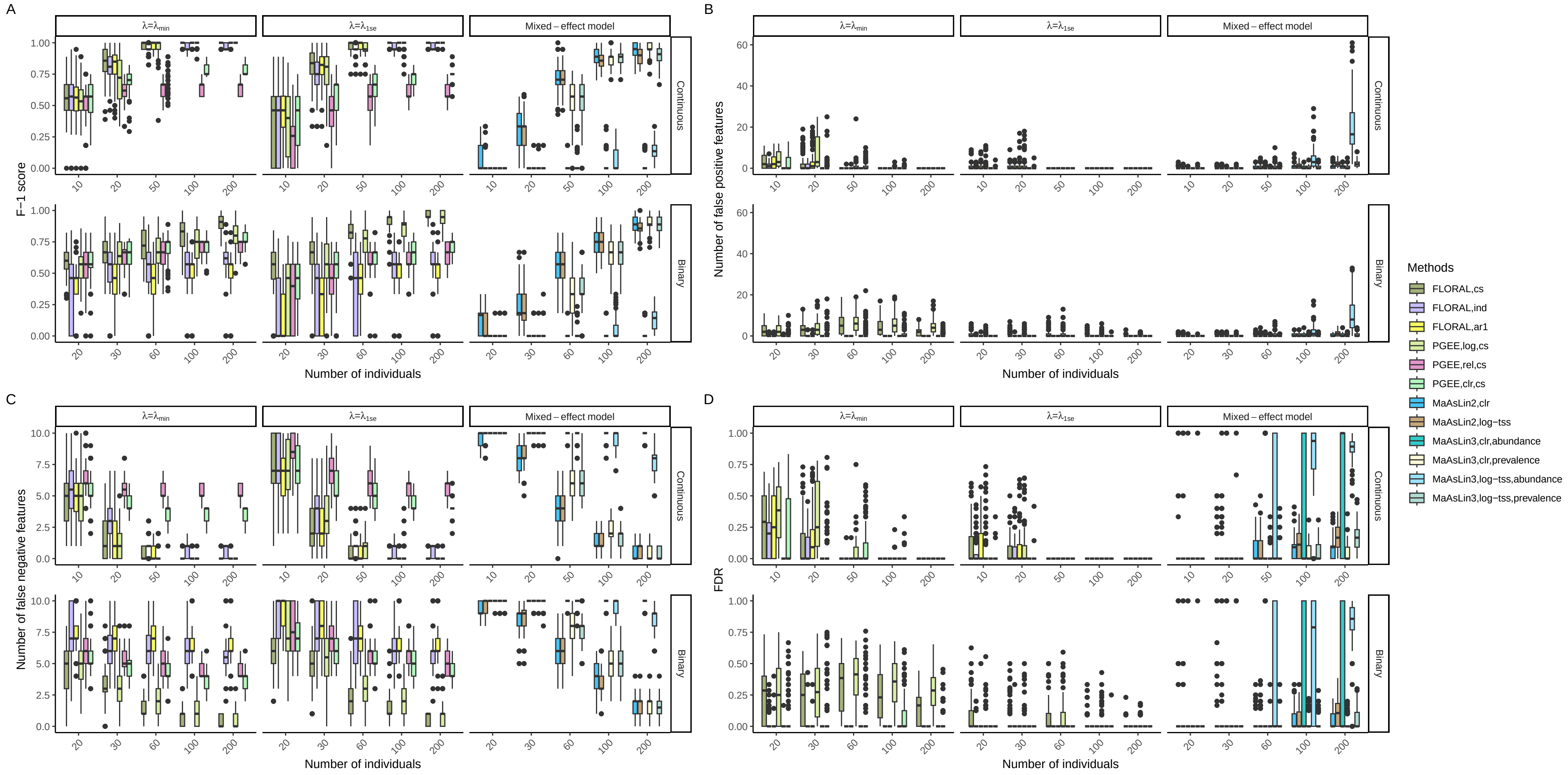


Fig. S1: Boxplots of **A.** $F_1$ score, **B.** number of false positive features, **C.** number of false negative features, and **D.** false discovery rate (FDR) obtained by PGEE and mixed-effect modeling methods for continuous and binary longitudinal with different number of individuals, where there were 10 true features in each simulation run. The reference scenario is $p = 200, m = 3, u = 0.15, \rho = 0, \kappa = 2.5$ for continuous outcome and $p = 200, m = 3, u = 0.3, \rho = 0, \kappa = 0$ for binary outcome.

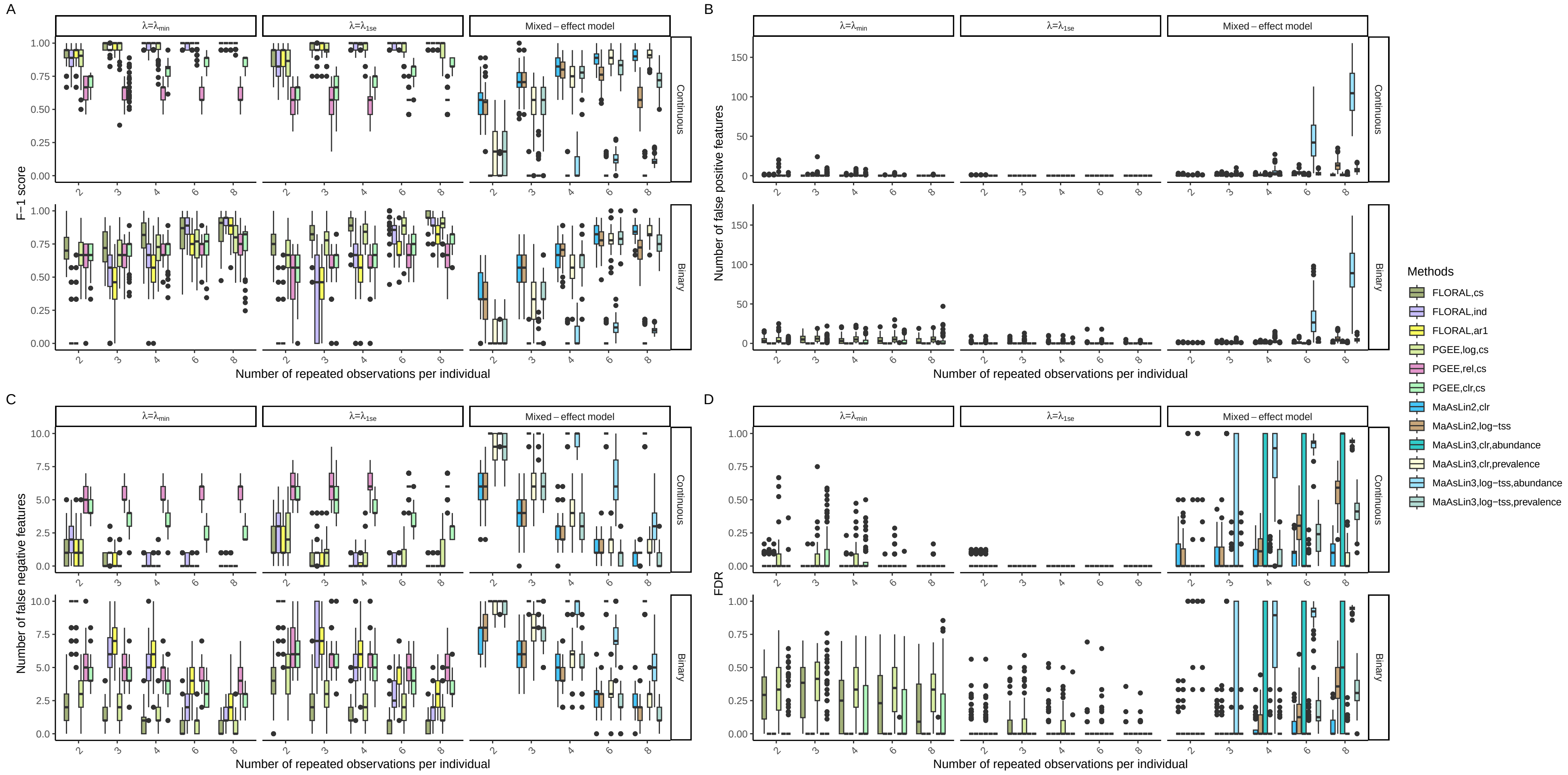


Fig. S2: Boxplots of **A.** $F_1$ score, **B.** number of false positive features, **C.** number of false negative features, and **D.** false discovery rate (FDR) obtained by PGEE and mixed-effect modeling methods for continuous and binary longitudinal with different number of repeated observations per individual, where there were 10 true features in each simulation run. The reference scenario is $n = 50, p = 200, u = 0.15, \rho = 0, \kappa = 2.5$ for continuous outcome and $n = 60, p = 200, u = 0.3, \rho = 0, \kappa = 0$ for binary outcome.

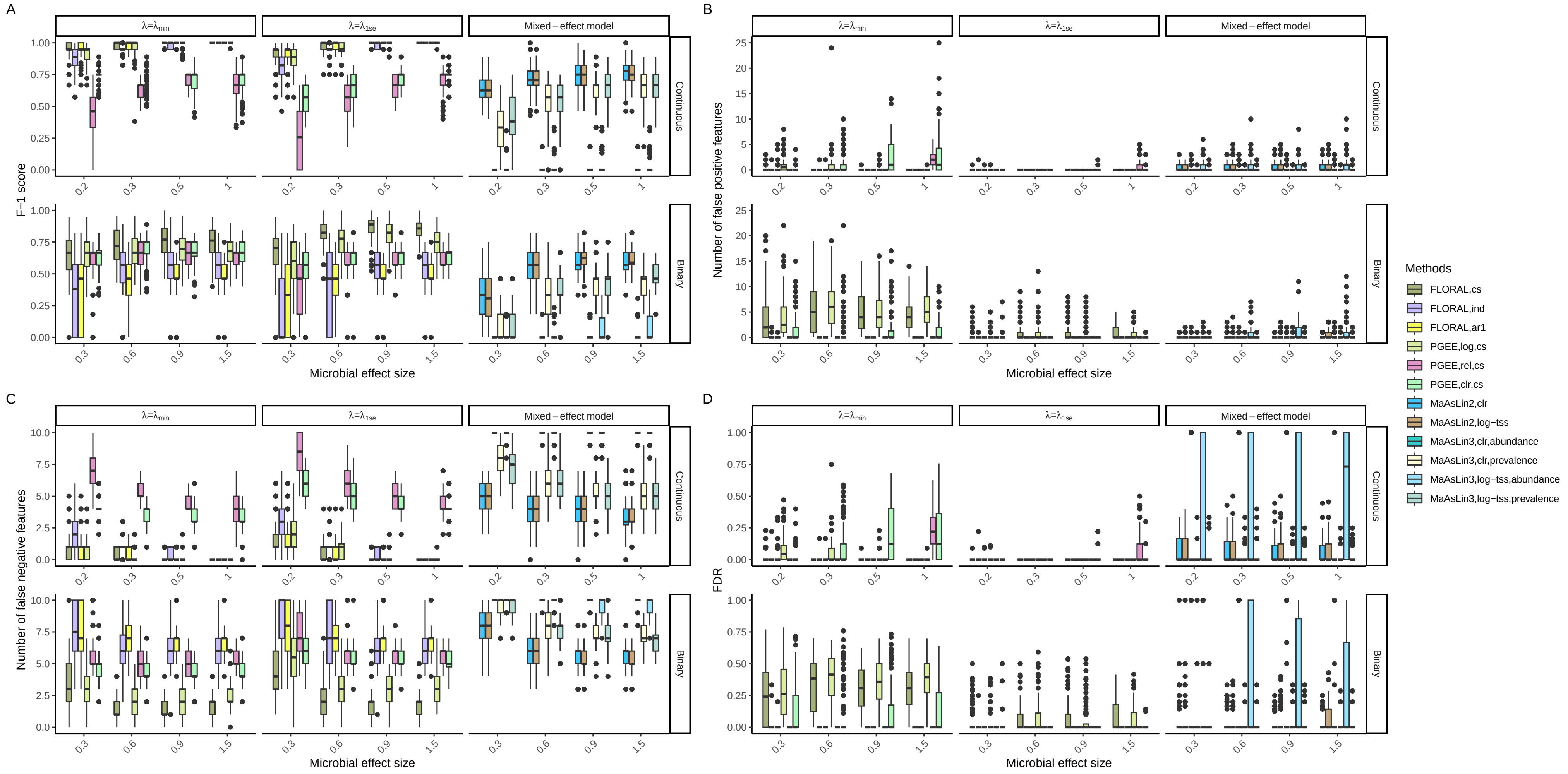


Fig. S3: Boxplots of **A.** $F_1$ score, **B.** number of false positive features, **C.** number of false negative features, and **D.** false discovery rate (FDR) obtained by PGEE and mixed-effect modeling methods for continuous and binary longitudinal with different microbial feature effect sizes, where there were 10 true features in each simulation run. The reference scenario is $n = 50, p = 200, m = 3, \rho = 0, \kappa = 2.5$ for continuous outcome and $n = 60, p = 200, m = 3, \rho = 0, \kappa = 0$ for binary outcome.

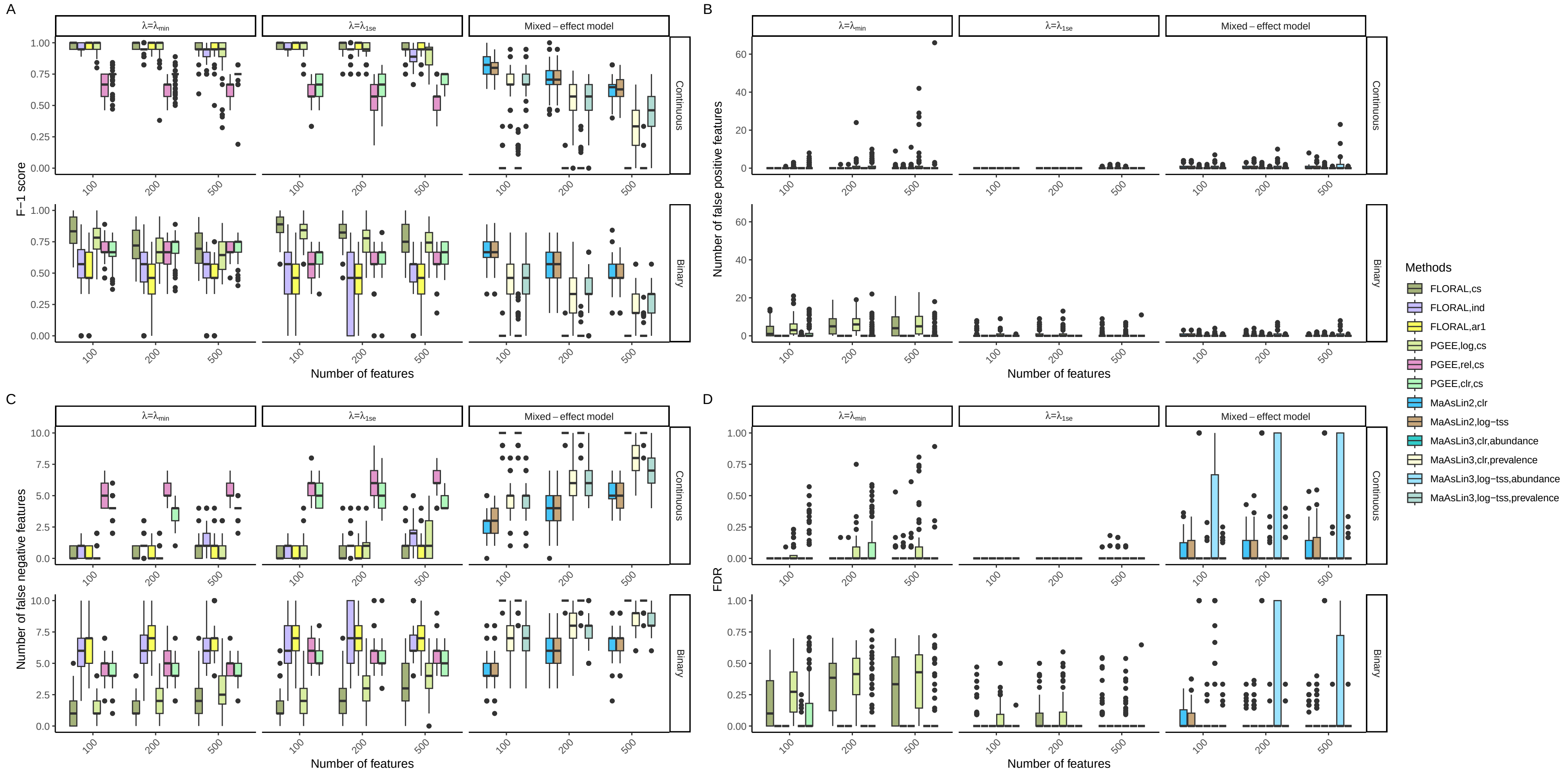


Fig. S4: Boxplots of **A.** $F_1$ score, **B.** number of false positive features, **C.** number of false negative features, and **D.** false discovery rate (FDR) obtained by PGEE and mixed-effect modeling methods for continuous and binary longitudinal with different number of microbial features, where there were 10 true features in each simulation run. The reference scenario is $n = 50, m = 3, u = 0.15, \rho = 0, \kappa = 2.5$ for continuous outcome and $n = 60, m = 3, u = 0.3, \rho = 0, \kappa = 0$ for binary outcome.

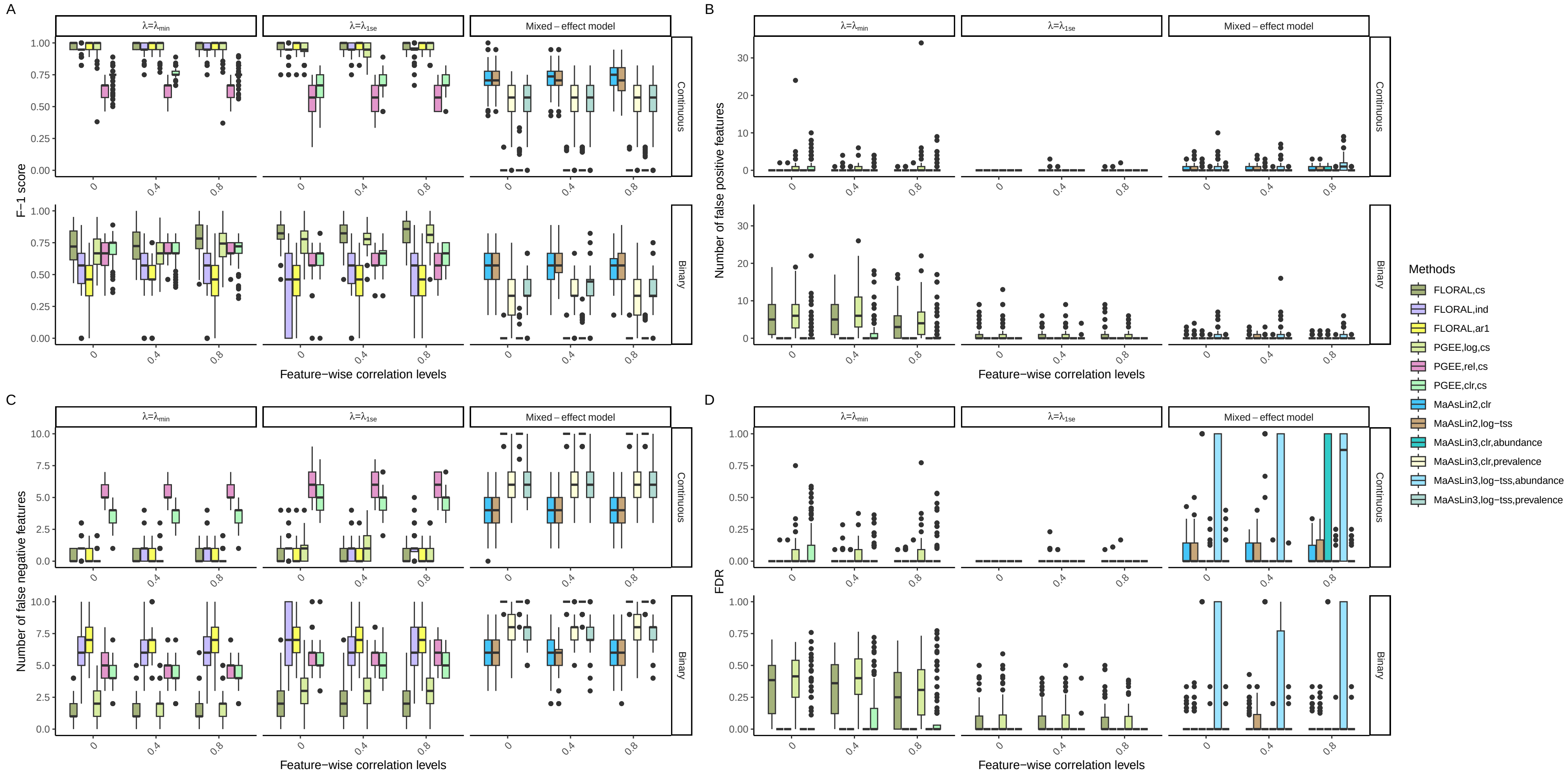


Fig. S5: Boxplots of **A.** $F_1$ score, **B.** number of false positive features, **C.** number of false negative features, and **D.** false discovery rate (FDR) obtained by PGEE and mixed-effect modeling methods for continuous and binary longitudinal with different feature-wise correlation levels, where there were 10 true features in each simulation run. The reference scenario is $n = 50, m = 3, p = 200, u = 0.15, \kappa = 2.5$ for continuous outcome and $n = 60, m = 3, p = 200, u = 0.3, \kappa = 0$ for binary outcome.

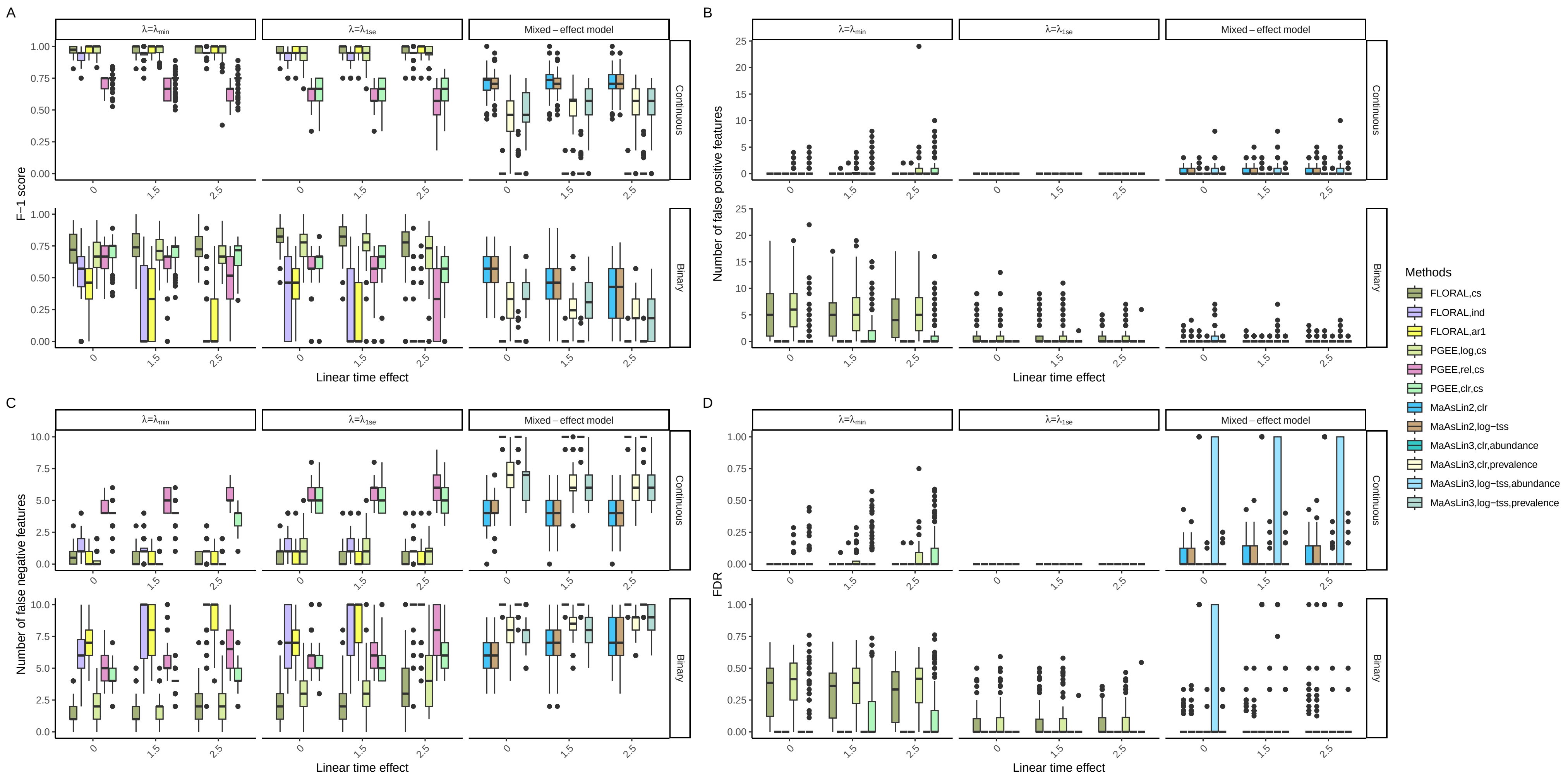


Fig. S6: Boxplots of **A.** $F_1$ score, **B.** number of false positive features, **C.** number of false negative features, and **D.** false discovery rate (FDR) obtained by PGEE and mixed-effect modeling methods for continuous and binary longitudinal with different linear time effect sizes, where there were 10 true features in each simulation run. The reference scenario is $n = 50, m = 3, p = 200, u = 0.15, \rho = 0$ for continuous outcome and $n = 60, m = 3, p = 200, u = 0.3, \rho = 0$ for binary outcome.

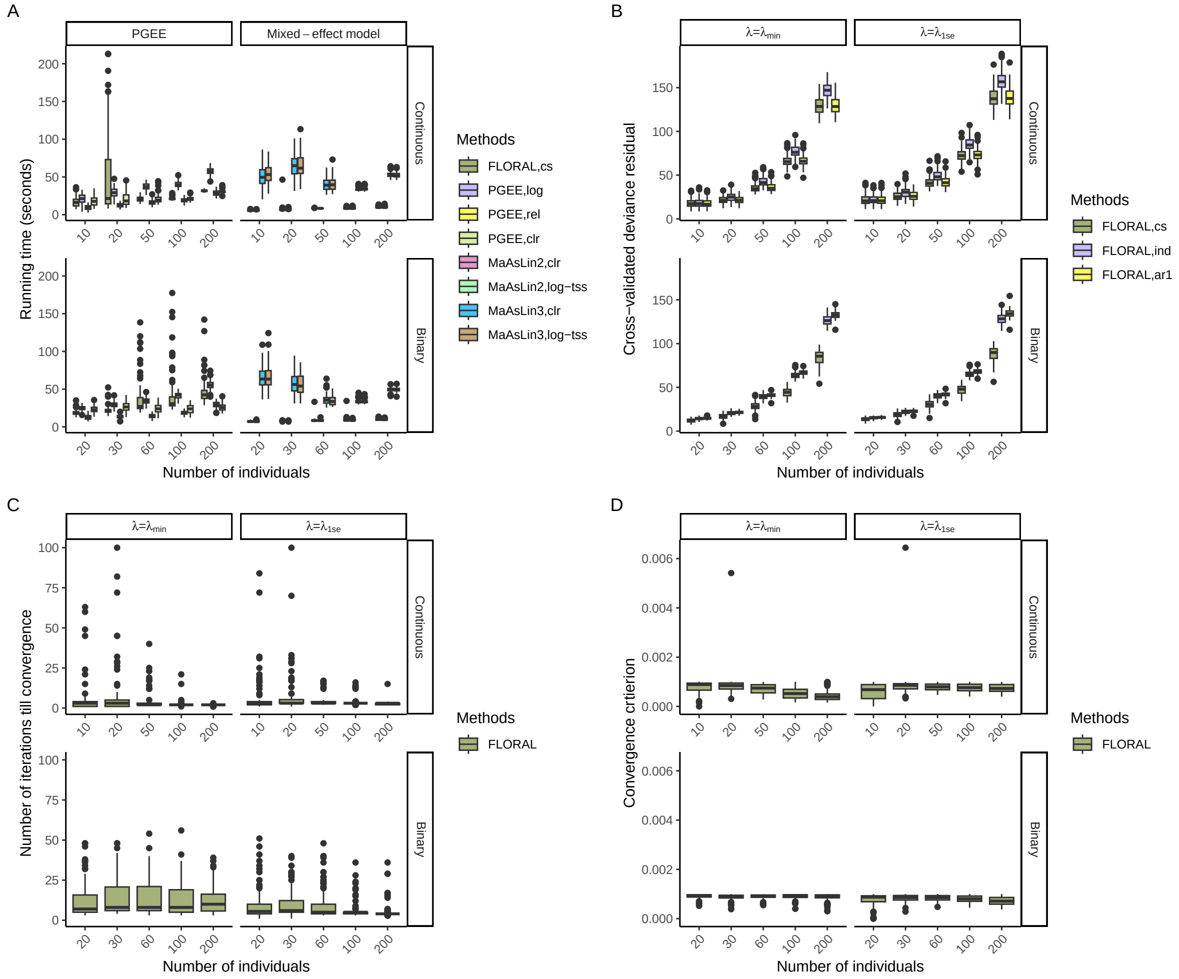


Fig. S7: Boxplots of **A.** running time in seconds of PGEE and mixed-effect models, **B.** cross-validated deviance residual obtained by `FLORAL` with different working correlation structures, **C.** number of iterations taken by `FLORAL` when the convergence was reached, and **D.** the value of convergence criterion when `FLORAL`'s algorithm stopped, as observed for 100 simulations with different number of individuals. The reference scenario is $p = 200, m = 3, u = 0.15, \rho = 0, \kappa = 2.5$ for continuous outcome and $p = 200, m = 3, u = 0.3, \rho = 0, \kappa = 0$ for binary outcome.

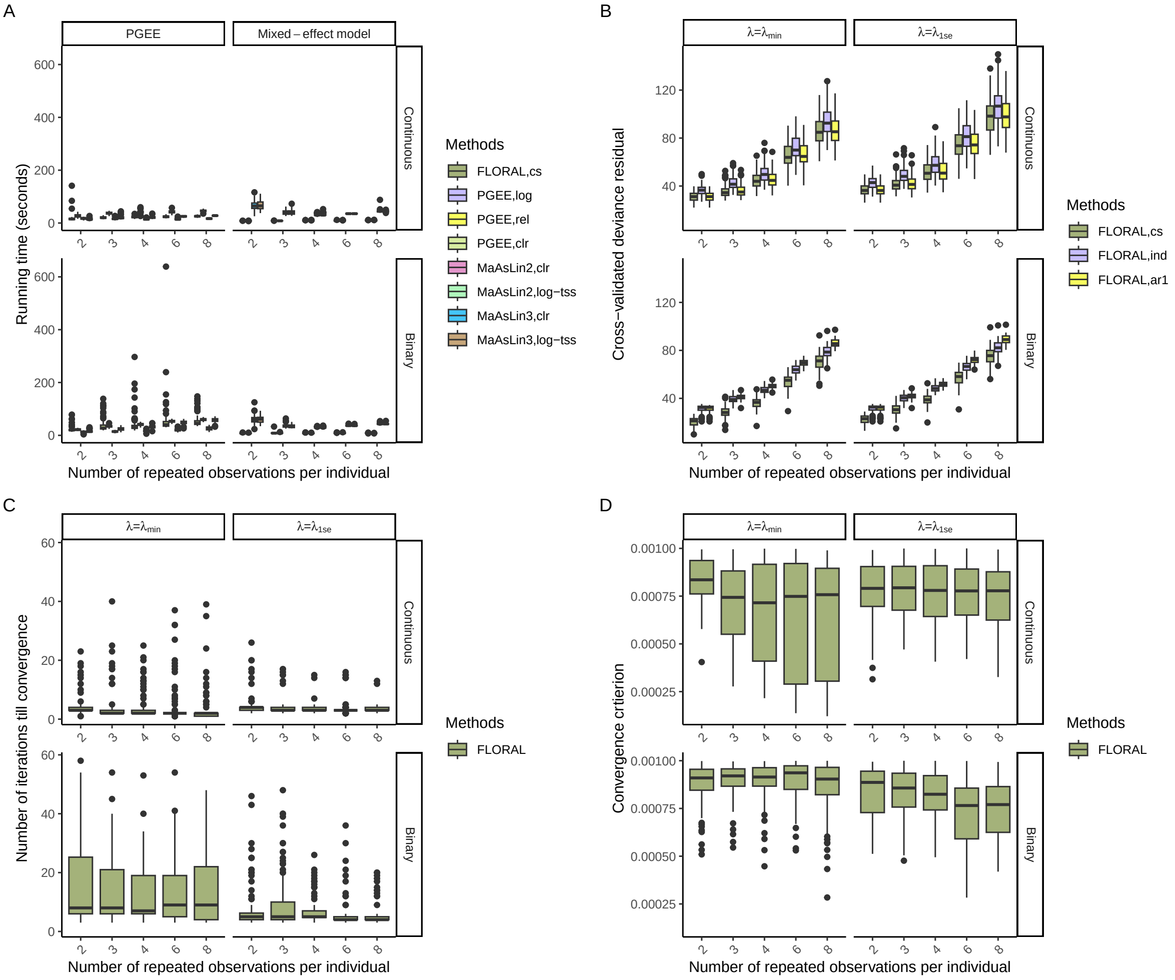


Fig. S8: Boxplots of **A.** running time in seconds of PGEE and mixed-effect models, **B.** cross-validated deviance residual obtained by `FLORAL` with different working correlation structures, **C.** number of iterations taken by `FLORAL` when the convergence was reached, and **D.** the value of convergence criterion when `FLORAL`'s algorithm stopped, as observed for 100 simulations with different number of repeated observations per individual. The reference scenario is $n = 50, p = 200, u = 0.15, \rho = 0, \kappa = 2.5$ for continuous outcome and $n = 60, p = 200, u = 0.3, \rho = 0, \kappa = 0$ for binary outcome.

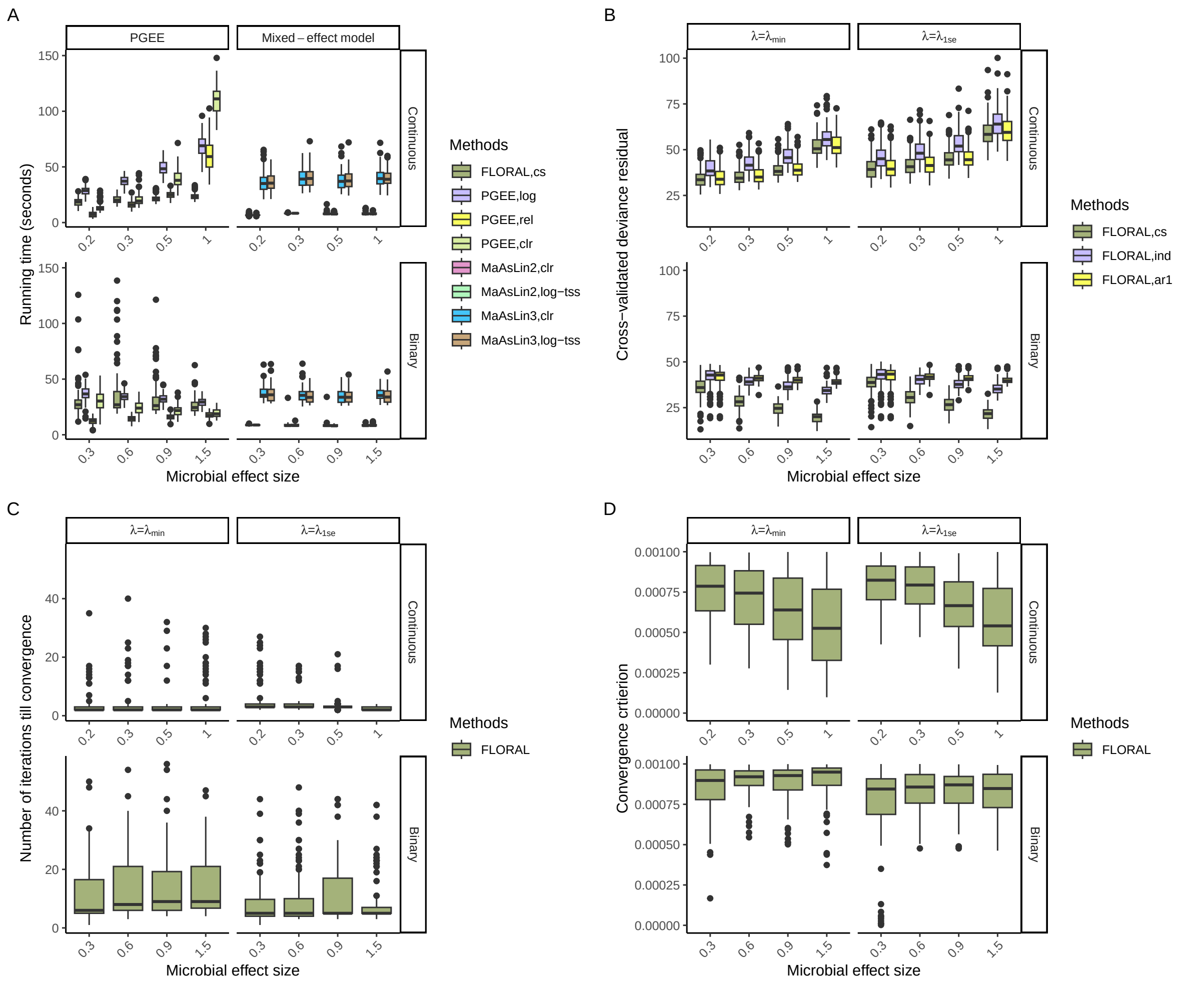


Fig. S9: Boxplots of **A.** running time in seconds of PGEE and mixed-effect models, **B.** cross-validated deviance residual obtained by `FLORAL` with different working correlation structures, **C.** number of iterations taken by `FLORAL` when the convergence was reached, and **D.** the value of convergence criterion when `FLORAL`'s algorithm stopped, as observed for 100 simulations with different microbial feature effect sizes. The reference scenario is $n = 50, p = 200, m = 3, \rho = 0, \kappa = 2.5$ for continuous outcome and $n = 60, p = 200, m = 3, \rho = 0, \kappa = 0$ for binary outcome.

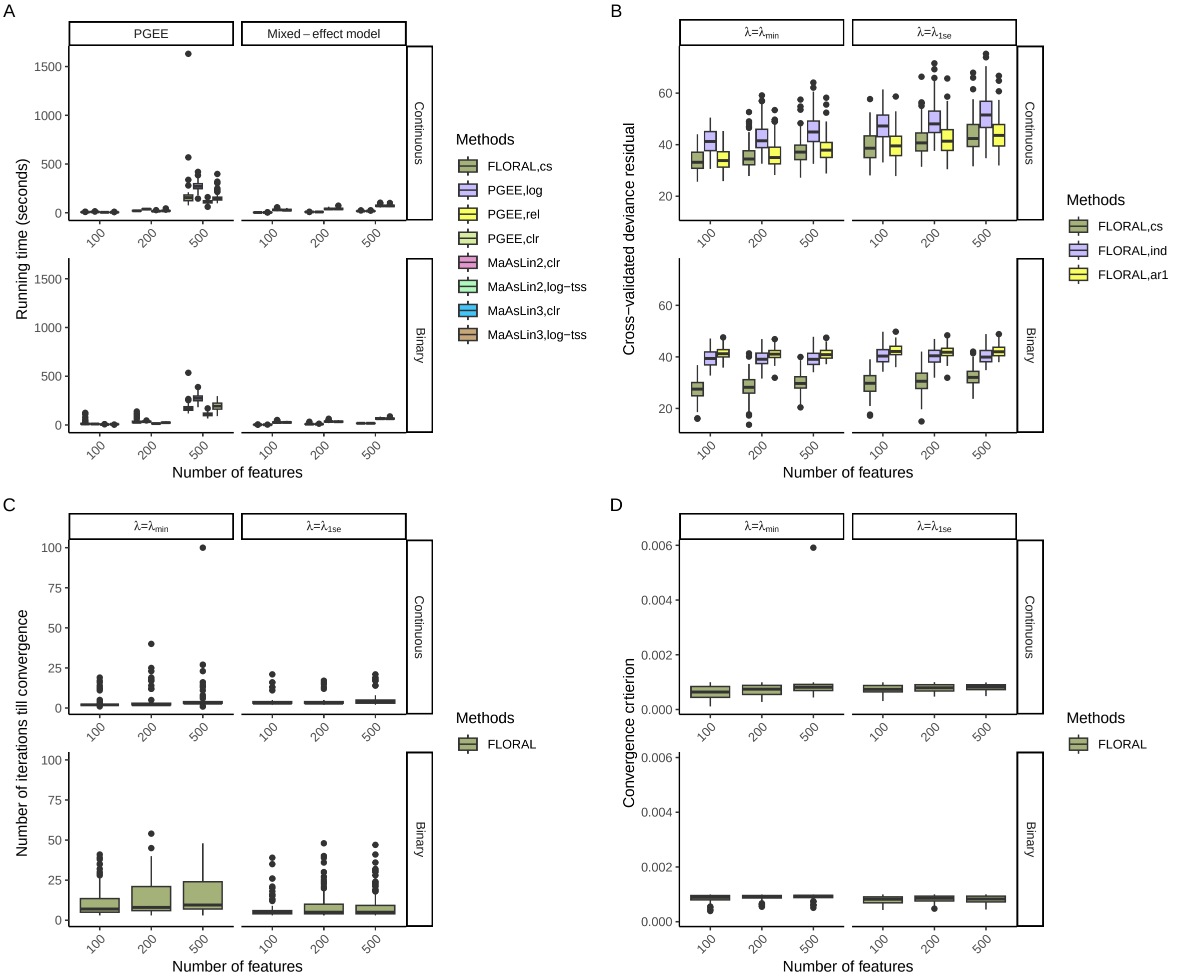


Fig. S10: Boxplots of **A.** running time in seconds of PGEE and mixed-effect models, **B.** cross-validated deviance residual obtained by `FLORAL` with different working correlation structures, **C.** number of iterations taken by `FLORAL` when the convergence was reached, and **D.** the value of convergence criterion when `FLORAL`'s algorithm stopped, as observed for 100 simulations with different number of microbial features. The reference scenario is $n = 50, m = 3, u = 0.15, \rho = 0, \kappa = 2.5$ for continuous outcome and $n = 60, m = 3, u = 0.3, \rho = 0, \kappa = 0$ for binary outcome.

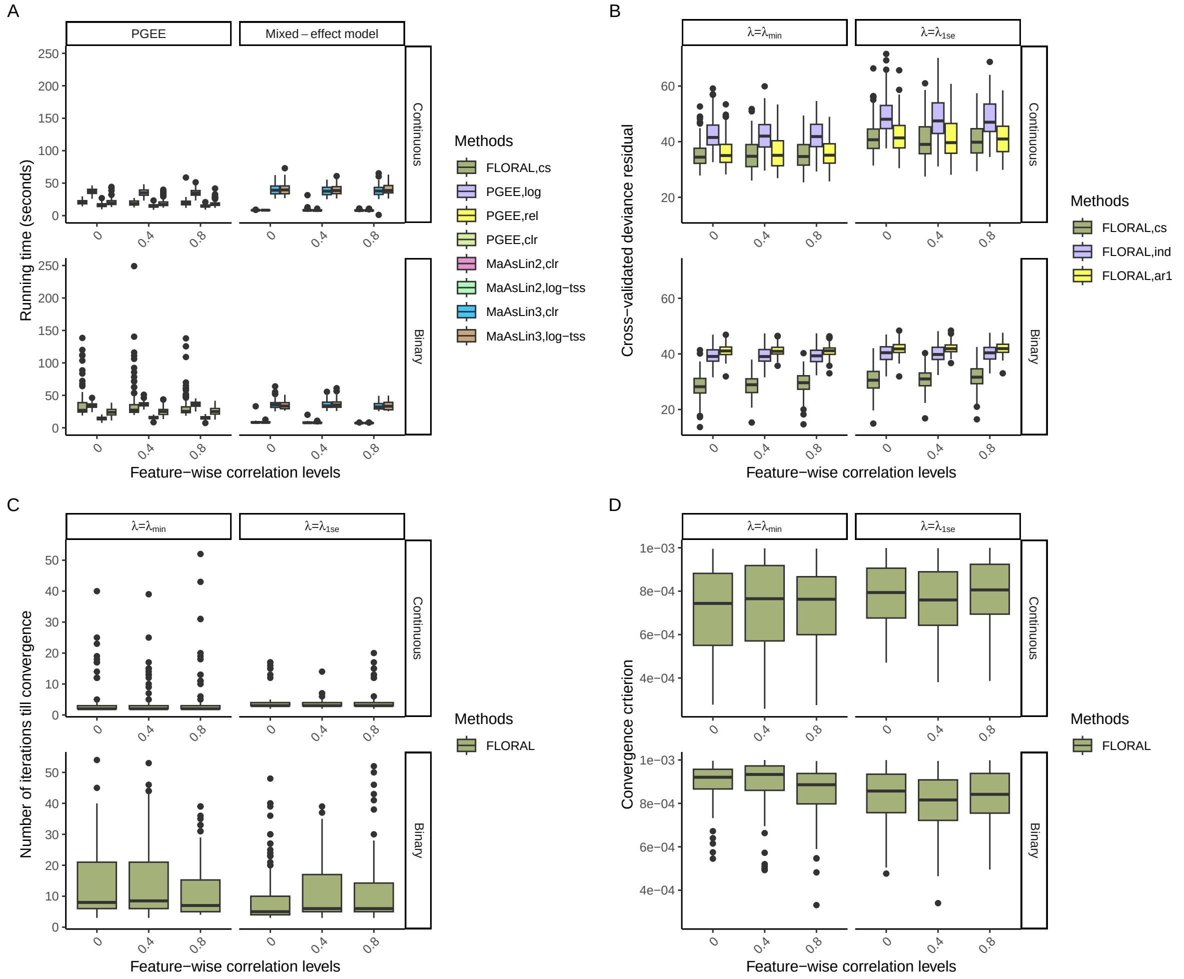


Fig. S11: Boxplots of **A.** running time in seconds of PGEE and mixed-effect models, **B.** cross-validated deviance residual obtained by `FLORAL` with different working correlation structures, **C.** number of iterations taken by `FLORAL` when the convergence was reached, and **D.** the value of convergence criterion when `FLORAL`'s algorithm stopped, as observed for 100 simulations with different feature-wise correlation levels. The reference scenario is $n = 50, m = 3, p = 200, u = 0.15, \kappa = 2.5$ for continuous outcome and $n = 60, m = 3, p = 200, u = 0.3, \kappa = 0$ for binary outcome.

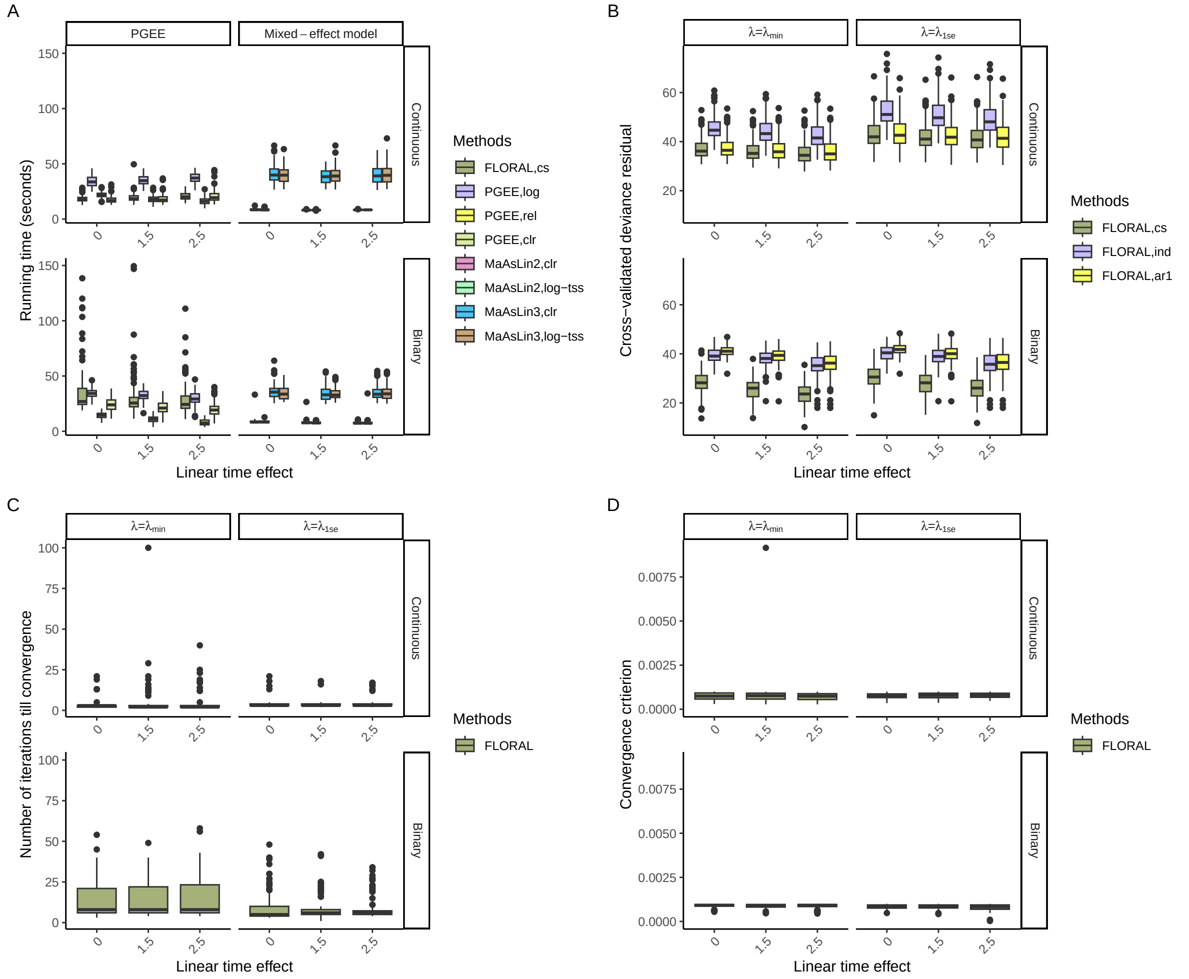


Fig. S12: Boxplots of **A.** running time in seconds of PGEE and mixed-effect models, **B.** cross-validated deviance residual obtained by `FLORAL` with different working correlation structures, **C.** number of iterations taken by `FLORAL` when the convergence was reached, and **D.** the value of convergence criterion when `FLORAL`'s algorithm stopped, as observed for 100 simulations with different linear time effect sizes. The reference scenario is $n = 50, m = 3, p = 200, u = 0.15, \rho = 0$ for continuous outcome and $n = 60, m = 3, p = 200, u = 0.3, \rho = 0$ for binary outcome.

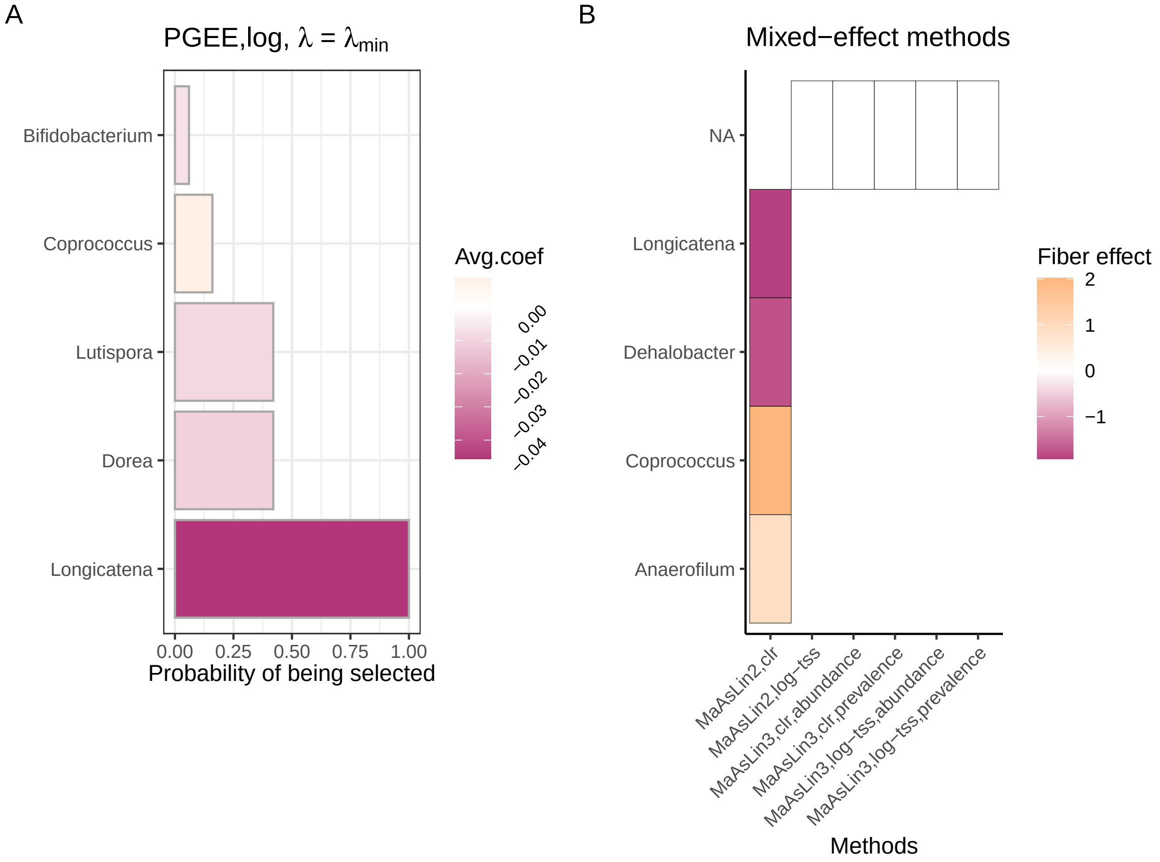


Fig. S13: Variable selection results for the NUTRIVENTION longitudinal fiber intake and microbiome data using the standard PGEE model and mixed-effect models `MaAsLin2` and `MaAsLin3`. **A.** Proportions of taxa being selected by `PGEE` with log-transformed data out of 100 runs of 5-fold cross-validation with random fold splits using compound symmetry structures. Colors represent the average feature coefficient out of 100 runs, where a positive coefficient implies a positive association between fiber intake and the microbial feature. **B.** Features selected by `MaAsLin2` and `MaAsLin3` with CLR-transformed (clr) and log-transformed total-sum scaling (log-tss) data. Selected features are marked by boxes with black boundaries. Colors represent the coefficients corresponding to the fiber effect on the abundance of microbial features. Methods detecting no features at the FDR level of 0.1 were visualized with 'NA'.

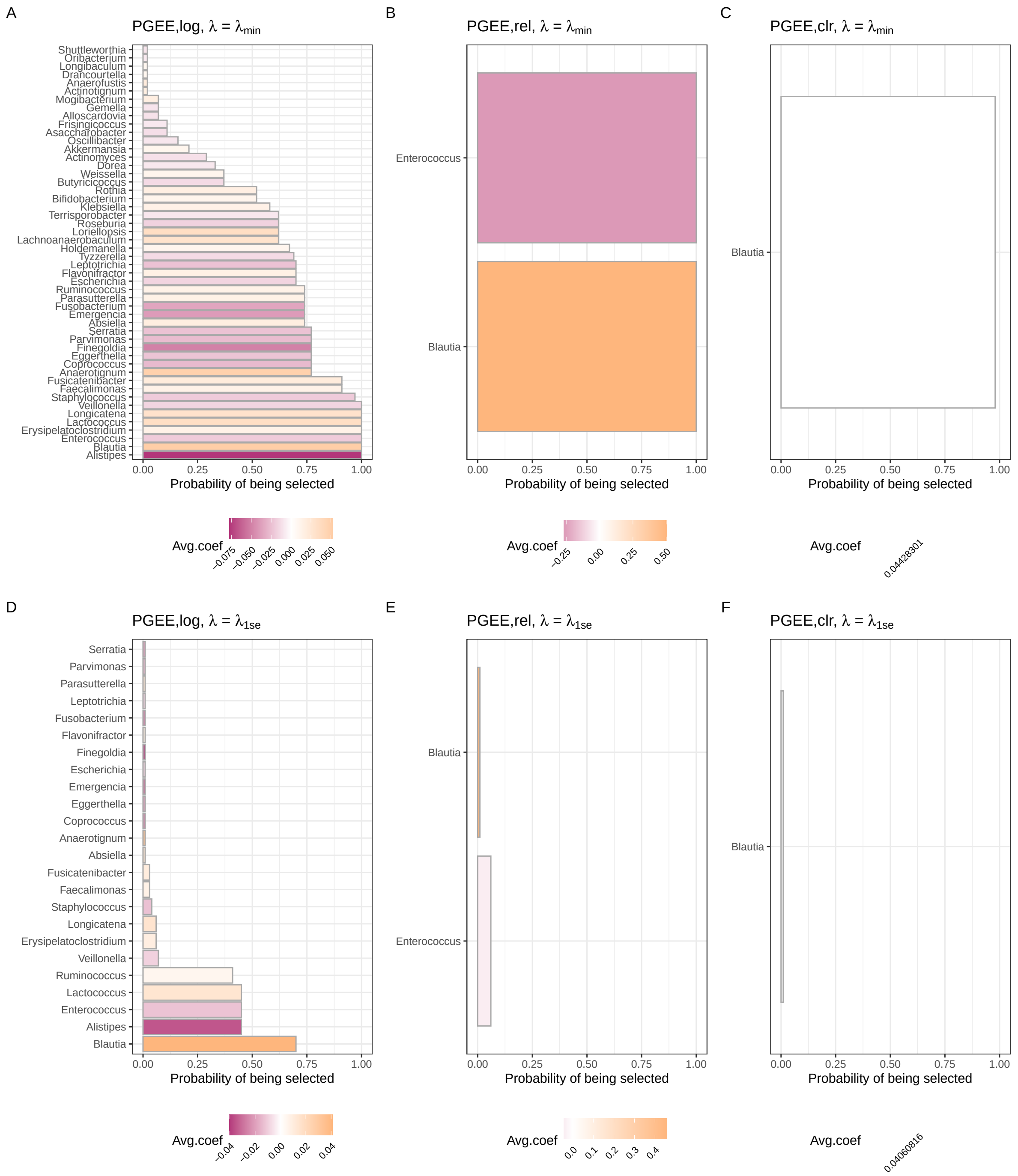


Fig. S14: Variable selection results for the allo-HCT longitudinal fiber intake and microbiome data using the standard PGEE method with $\lambda = \lambda_{\text{min}}$ (**A-C**) and $\lambda = \lambda_{\text{1se}}$ (**D-F**). **A-F.** Proportions of taxa being selected by `PGEE` with **A,D.** log-transformed (log), **B,E.** relative abundance (rel) and **C,F.** CLR-transformed (clr) data out of 100 runs of 5-fold cross-validation with random fold splits using compound symmetry structures. Colors represent the average feature coefficient out of 100 runs, where a positive coefficient implies a positive association between fiber intake and the microbial feature.

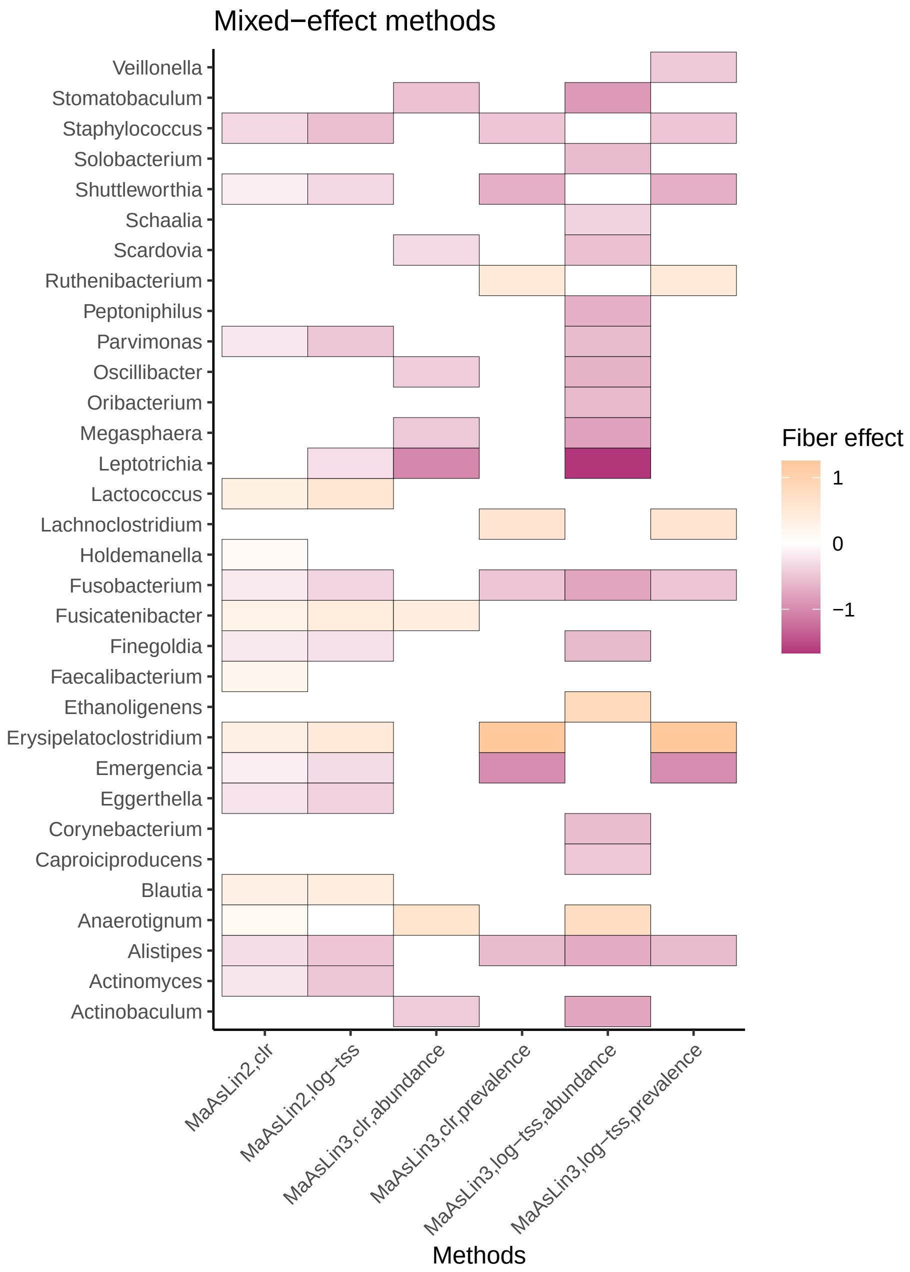


Fig. S15: Variable selection results for the allo-HCT longitudinal fiber intake and microbiome data at the FDR level of 0.1 using mixed-effect models `MaAsLin2` and `MaAsLin3` with different data transformation and normalization strategies, including CLR-transformation (clr) and log-transformed total-sum scaling (log-tss). Selected features are marked by boxes with black boundaries. Colors represent the coefficients corresponding to the fiber effect on the abundance of microbial features.

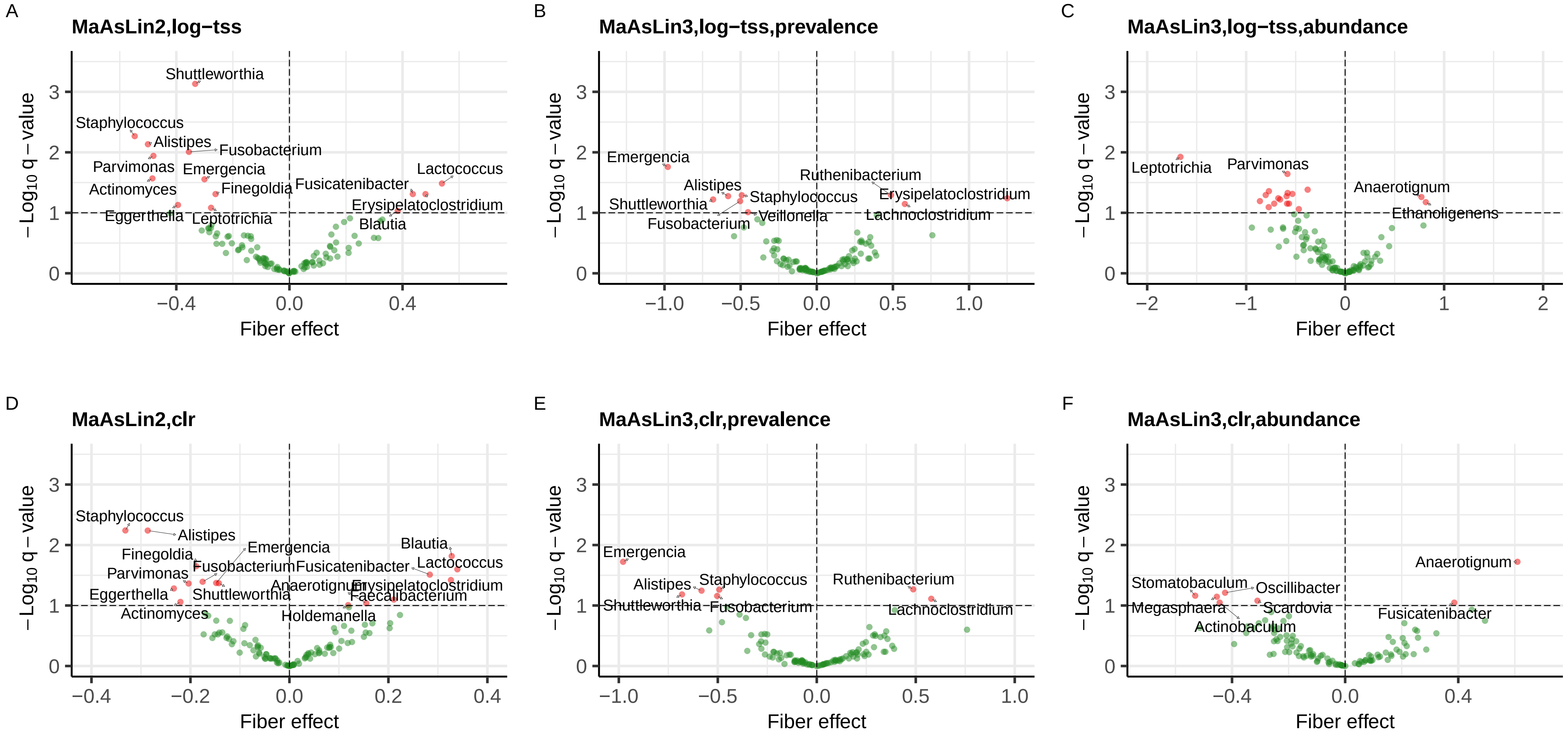


Fig. S16: Volcano plots showing the fiber intake effect size on microbial abundances in the allo-HCT cohort and the corresponding negative log-transformed q-values for `MaAsLin2` with **A.** log-transformed total-sum scaling (log-tss) and **D.** CLR-transformation (clr), `MaAsLin3` prevalence models with **B.** log-tss normalization and **E.** CLR-transformation, and `MaAsLin3` abundance models with **C.** log-tss normalization and **F.** CLR-transformation. Genera with the most significant associations with fiber intake are printed on the plots. The horizontal dashed line displays the q-value threshold of 0.1, while the vertical dashed line shows the zero fiber intake effect.

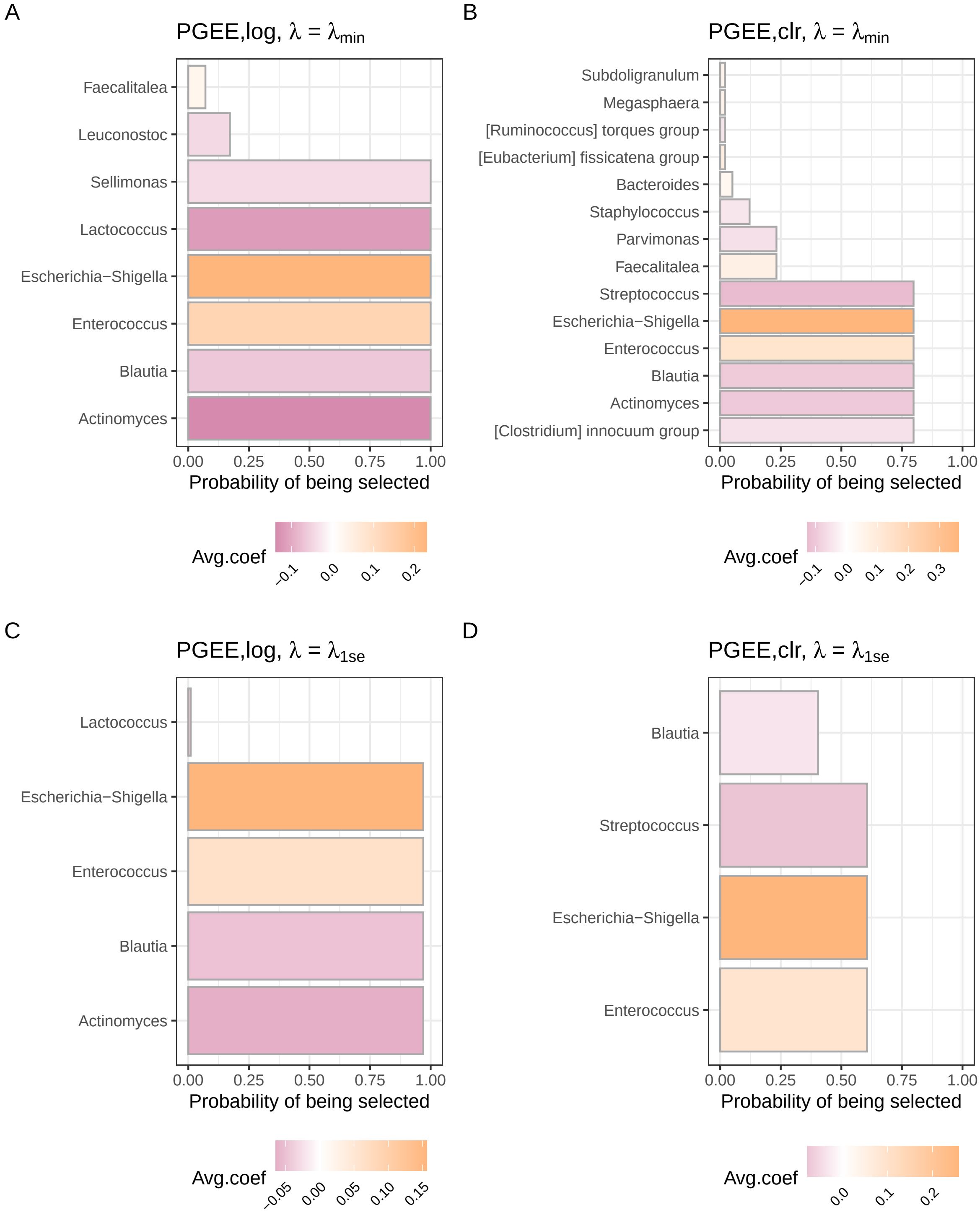


Fig. S17: Model fitting and variable selection results for the allo-HCT longitudinal BSI status and microbiome data using using the standard PGEE method with $\lambda = \lambda_{\text{min}}$ (**A-B**) and $\lambda = \lambda_{\text{1se}}$ (**C-D**). **A-D.** Proportions of taxa being selected by `PGEE` with **A,C.** log-transformed (log), **B,D.** CLR-transformed (clr) data out of 100 runs of 5-fold cross-validation with random fold splits using compound symmetry structures. Colors represent the average feature coefficient out of 100 runs, where a positive coefficient implies a positive association between a positive BSI status and the microbial feature.

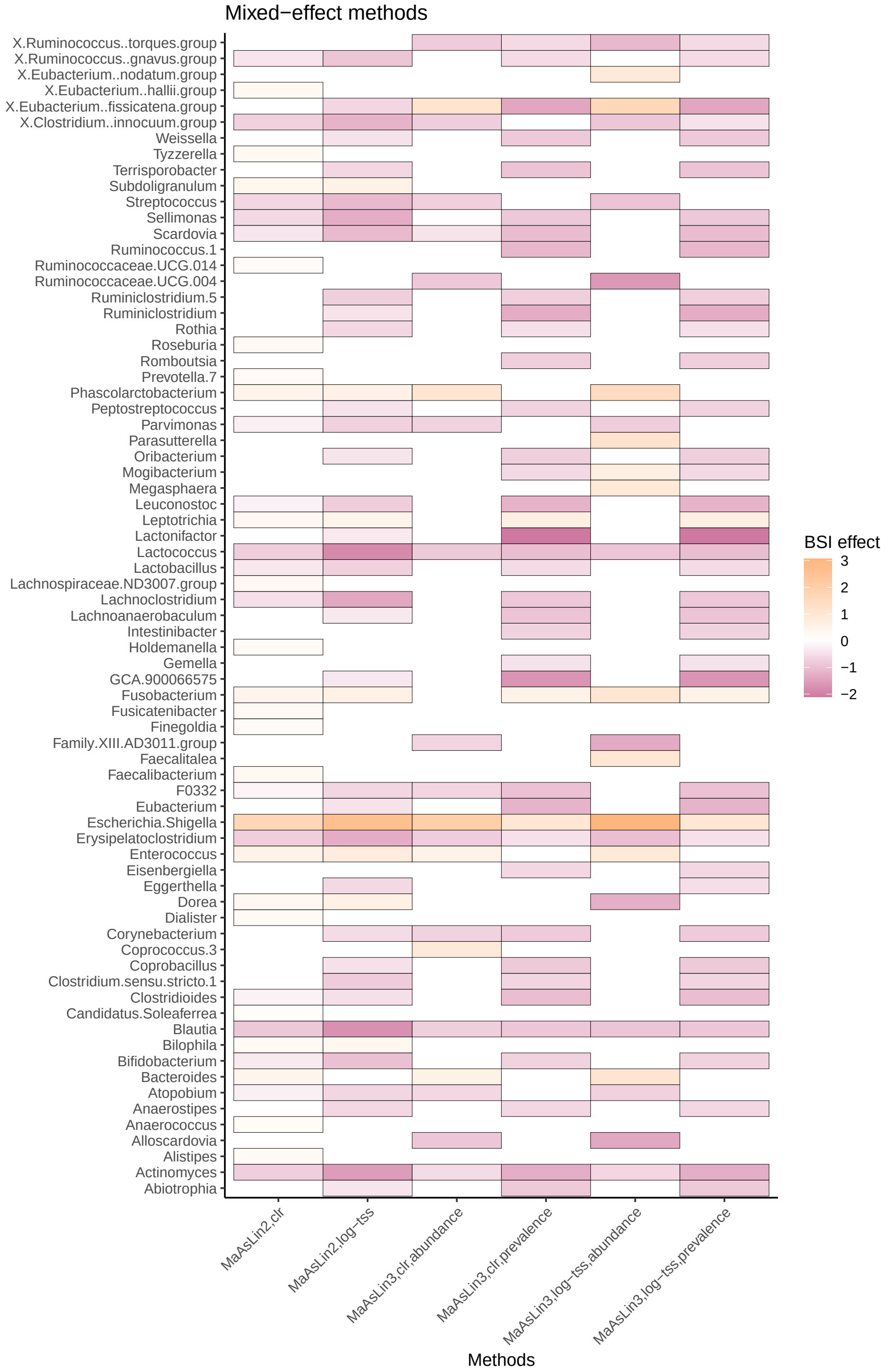


Fig. S18: Variable selection results for the allo-HCT longitudinal BSI status and microbiome data at the FDR level of 0.1 using mixed-effect models `MaAsLin2` and `MaAsLin3` with different data transformation and normalization strategies, including CLR-transformation (clr) and log-transformed total-sum scaling (log-tss). Selected features are marked by boxes with black boundaries. Colors represent the coefficients corresponding to the fiber effect on the abundance of microbial features.

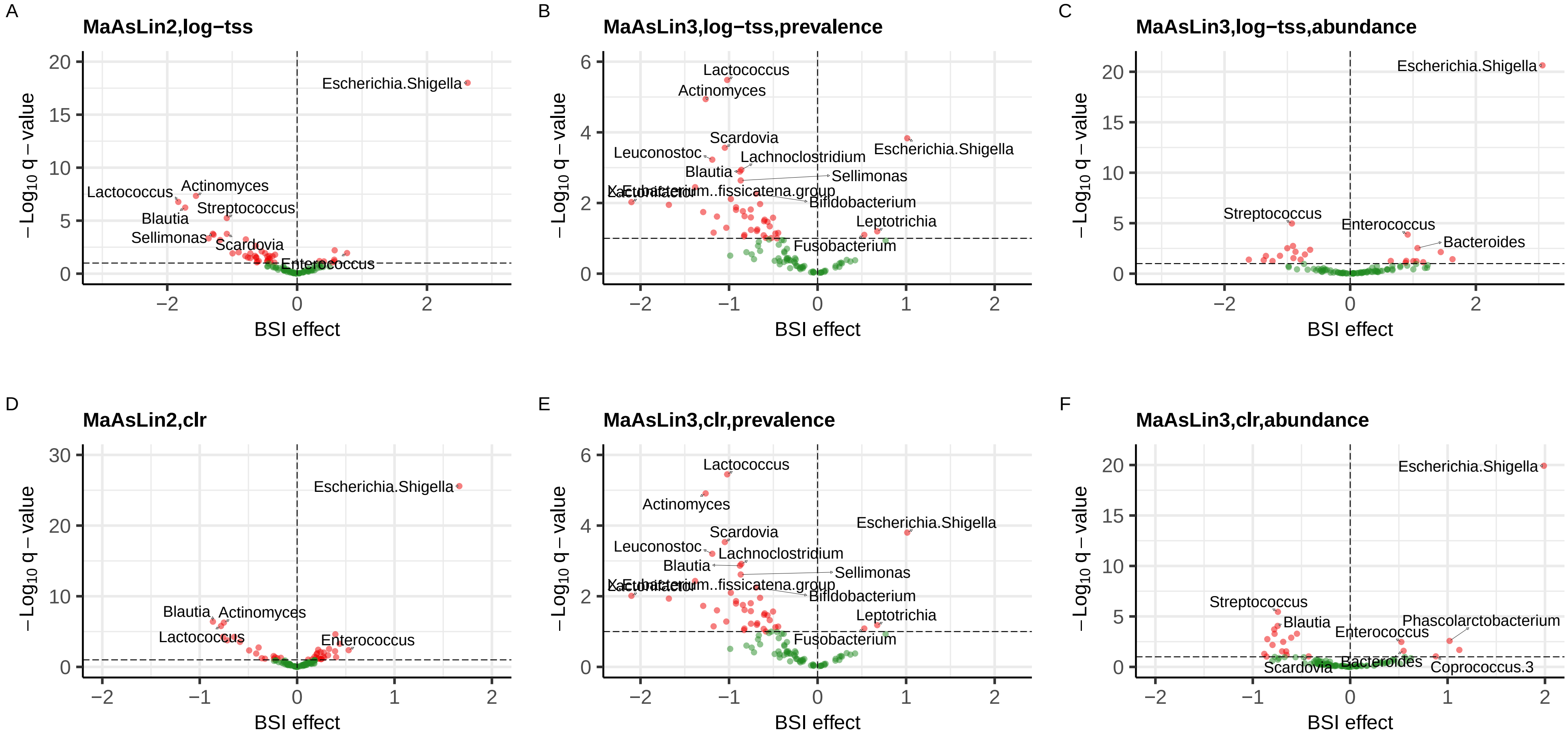


Fig. S19: Volcano plots showing the BSI status effect size on microbial abundances in the allo-HCT cohort and the corresponding negative log-transformed q-values for `MaAsLin2` with **A.** log-transformed total-sum scaling (log-tss) and **D.** CLR-transformation (clr), `MaAsLin3` prevalence models with **B.** log-tss normalization and **E.** CLR-transformation, and `MaAsLin3` abundance models with **C.** log-tss normalization and **F.** CLR-transformation. Genera with the most significant associations with fiber intake are printed on the plots. The horizontal dashed line displays the q-value threshold of 0.1, while the vertical dashed line shows the zero fiber intake effect.
